# The BEAC, an epigenetic clock for birds

**DOI:** 10.64898/2026.08.14.744821

**Authors:** Mikaela Hukkanen, Simon Jarman, Alyssa Budd, Flávia Akemi Nitta Fernandes, Roberto Ambrosini, Chloe Anderson, Gaël Bardon, Oliver Berry, Pierre-Paul Bitton, Thomas Bugnyar, Manuela Caprioli, Nicholas Carlile, Jacopo G. Cecere, Nina Cossin-Sevrin, Alessandra Costanzo, Alejandro Corregidor-Castro, Leyla Rivero Davis, Eva van Dijk, Michelle Elsner, Kyle H. Elliott, Joan Ferrer Obiol, Didone Frigerio, Nino Gardoni, Philippe Helsen, Maureen Hofer, Sonia Kleindorfer, Joost Lammers, Don-Jean Léandri-Breton, Jorg Massen, Guillam E. McIvor, Britta S. Meyer, Antoine Morel, Michelangelo Morganti, Elodie Paciello, Josephine Paris, Andrea Pilastro, Leonie Pihlflyckt, Paula Plaza, Andrea M. Polanowski, Aada Puhakka, Lauren Roman, Andrea Romano, Diego Rubolini, Akshata Shetty, Petra Sumasgutner, Conor Taff, Alex Thornton, Emiliano Trucchi, Maren Vitousek, Sofia Westerlund, Shannon Whelan, Sandrine Zahn, Jaakko Kaprio, Céline Le Bohec, Miina Ollikainen, Robin Cristofari

## Abstract

Epigenetic clocks are powerful tools for estimating both chronological and biological age, enabling the integration of age information into population monitoring, demographic modelling, and research on the ecophysiology and evolution of ageing. Most epigenetic clocks so far have been developed for mammals: here, we present the Bird Epigenetic Ageing Clock (BEAC) for estimating chronological age in avian species. BEAC was established based on genome-wide enzymatic methylation sequencing data of known-age king penguins (*Aptenodytes patagonicus*), and validated in nine other bird species. The BEAC collects age-informative signals into a bisulfite amplicon sequencing panel of 24 primer pairs, providing a highly accurate and cost-effective alternative to sequencing-intensive approaches. It achieved strong predictive performance in independent king penguin training (R²=0.88; MAE=1.7 years, n=78) and testing data (R²=0.79; MAE=2.3 years, n=41), with negligible batch effects, high longitudinal consistency, and resilience to reduced sample size or missing loci. Importantly, cross-species validation across 180 samples showed that BEAC reliably captures age-associated methylation signals in nine additional bird species across seven clades, demonstrating that a single set of loci can be predictive of ageing across multiple different bird species. BEAC offers a flexible, empirically validated tool and a transferable framework for developing epigenetic clocks in avian species, providing a highly valuable resource for eco-evolutionary studies of ageing in wild species.

## Introduction

Epigenetic clocks are computational algorithms based on methylation levels at the fifth carbon of a cytosine base, in the context of a CG dinucleotide (CpG-site)^1^. These DNA methylation-based algorithms are currently the most powerful biomarkers for predicting an individual’s age, with significant associations with morbidity and mortality in humans^2,3^ and utility across a range of other applications, from population viability analyses for endangered species^4,5^ and management of commercial animals^6,7^ to geroscience and the treatment of age-related diseases^8^. Epigenetic clocks have also been used to infer the genetic and environmental drivers of ageing rate both in humans and in wild animals^9–13^. Most existing clocks rely on microarrays designed specifically for humans or other mammals^14–18^, thus limiting their practical applications and, importantly, the biological understanding of ageing in phylogenetically diverse taxa.

The development of epigenetic clocks for non-mammalian taxa is currently constrained by a trade-off between genomic resolution and utility at scale. To date, many non-mammalian clocks have relied on inferring methylation levels using reduced representation techniques such as RRBS^13,23,24^, bsRAD-seq^25–27^, and DREAM^28^. While these bypass the need for species-specific microarrays, they only capture methylation at selected restriction enzyme sites, potentially overlooking the most age-informative CpG-sites located elsewhere in the genome^29–31^. Furthermore, as sample size increases, “locus dropout” occurs due to (i) restriction enzyme cut variability, (ii) PCR stochasticity, (iii) library complexity limits, (iv) mapping differences, that lead to a shrinking set of shared loci across individuals, making it difficult to merge datasets or replicate findings^29–31^. A few studies have used whole-genome methylation sequencing^32,33^, offering an unbiased view of the entire methylome, but it may be cost-prohibitive for the large cohorts required in ecological and population-level research^31^. To bridge this gap, some researchers^23,34,35^ have transitioned to targeted enrichment assays for validated age-related sites, such as multiplexed bisulfite amplicon sequencing^36^, significantly reducing per-sample costs. However, these remain rare and are still tethered to the original “biased” discovery data (a priori selected genes^35,37^, RRBS^23^ or ribosomal DNA^34^). In the context of avian biology, this gap is particularly pronounced: to our knowledge, there is no validated and cost-effective assay that would allow methylation typing across multiple bird species, leaving a significant void in the ornithological toolkit.

Mammalian research has built a robust foundation for our understanding of epigenetic ageing, but the sheer diversity of life histories across the vertebrate tree makes it difficult to determine if the genomic bases for these mechanisms are truly conserved. Birds diverged from the mammalian lineage over 300 million years ago^38^, and thus offer a unique lens through which to view the evolutionary stability of epigenetic markers. Pioneering studies have already developed successful models for species such as short-tailed shearwater (*Ardenna tenuirostris*)^28^, domestic chicken (*Gallus gallus domesticus*)^33^, great tit (*Parus major*)^24^ and chestnut-crowned babbler (*Pomatostomus ruficeps*)^39^. These foundational works, particularly those utilizing whole-genome methylation sequencing^33,39^, have demonstrated that age-related DNA methylation is a quantifiable avian trait. However, it remains unclear whether these patterns can be extended to build epigenetic ageing markers for use across multiple bird species. Birds have diverse life histories and lifespans that vary from three years in grassbirds to more than 80 years in cockatoos^40^, yet, whether ageing markers are conserved across species with different lifespans remains unresolved.

In this study, we describe the BEAC (Bird Epigenetic Ageing Clock) – an epigenetic clock designed from whole-genome methylation data for the long-lived king penguin (*Aptenodytes patagonicus*), a seabird with maximum lifespan of at least 30 years in the wild^41^, and 43 years in human care^41^. The king penguin represents a particularly valuable system for epigenetic clock development because decades of monitoring at the Crozet Islands have produced an exceptionally well-characterized population of known-age individuals^42^, overcoming one of the major limitations in developing ageing biomarkers for wild species^16^. Although initially targeted for king penguins, we demonstrate broader application of this clock across seven neognath orders. To improve access to targeted epigenetic ageing tools for non-model taxa, we describe BEAC-Seq, a multiplexed bisulfite amplicon sequencing protocol targeting age-associated methylation sites, and the BEAC-Model, an accompanying prediction algorithm. Together, these tools provide a cost-effective and scalable alternative to genome-wide sequencing for avian species. The BEAC-Model is calibrated employing elastic-net machine-learning^43^ on a wild population of king penguins, while the BEAC-Seq is tested on 14 bird species, of which 12 had sufficient sample sizes for age-association analyses. Together, these tools facilitate epigenetic age estimation in non-model bird species and provide a framework for testing whether age-associated methylation patterns are conserved across avian taxa with diverse life histories.

## Results

### BEAC-Seq design

We developed a bisulfite amplicon sequencing protocol^36^, the Bird Epigenetic Ageing Clock sequencing assay (hereafter BEAC-Seq), that allows targeted amplification and methylation typing at 24 loci, allowing the calculation of epigenetic age without the need for whole genome methylation sequencing. Briefly, we first used whole-genome Enzymatic Methylation Sequencing^44^ (EM-Seq) from 96 whole-peripheral-blood DNA samples from 87 unrelated known-age king penguins for marker discovery (ages 0.88–23.12 years, Figure 1, Supplementary Table 1, Methods). Because age-related methylation changes may follow either linear or non-linear monotonic trends, we screened all interrogated CpG-sites using both Pearson and Spearman correlations with age. The 100 most significant CpG-sites from each analysis were retained, resulting in a combined set of 118 unique candidate CpG-sites for downstream assay design. Targeting the selected CpG-sites, we designed flanking primer pairs using the king penguin reference genome (GCA_010087175.1, used throughout the manuscript)^45^ as a template. Primer pairs were designed with the following core constraints: (1) exclusion of CpG-sites in repetitive regions for higher specificity, (2) avoidance of known Single Nucleotide Polymorphisms (SNPs) to ensure unbiased amplification, (3) avoidance of CpG dinucleotides within the primer binding regions and (4) prioritization of conserved sequences to facilitate broad-scale application across bird species^46^. The resulting BEAC-Seq assay consists of 24 primer pairs covering 82 CpG-sites associated with 23 distinct genomic regions (Supplementary Table 2), with amplicon sizes ranging from 100 to 147 basepairs (bp). Figure 2 illustrates the methylation dynamics of these targeted sites, showing the relationship of methylation and age for one CpG per amplicon, selected with the highest adjusted R^2^ with log(Age).

**Figure 1.**
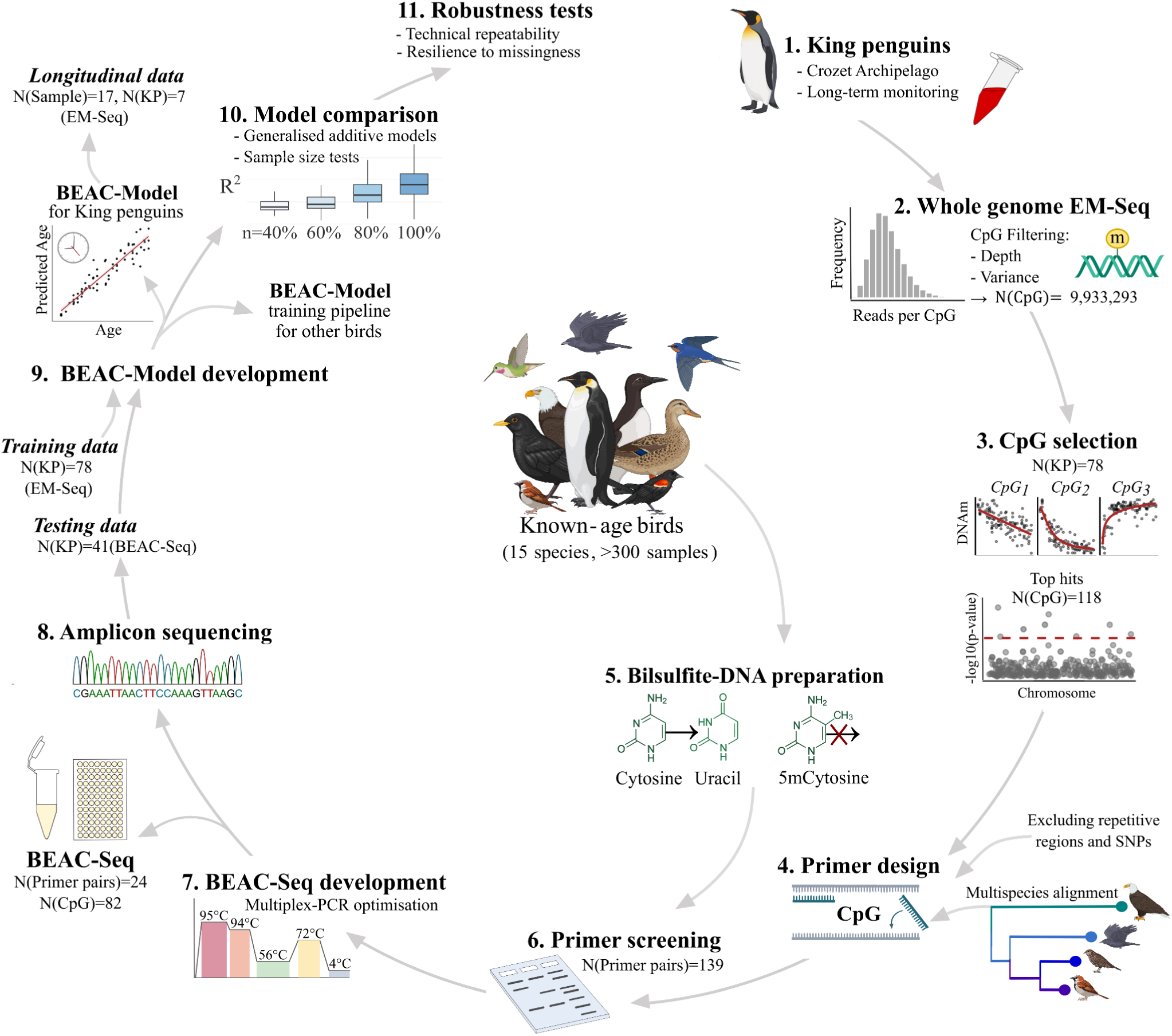
Flowchart of steps included in the development of the Bird Epigenetic Ageing Clock (BEAC). Steps 1-11 detail the identification of age-informative CpG sites from king penguin (KP) enzymatic sequencing (EM-Seq) data; design of the targeted bisulfite amplicon sequencing protocol; and clock training and performance validation across missing data scenarios, technical replicates, and additional bird species. Figure created in BioRender.com

**Figure 2.**
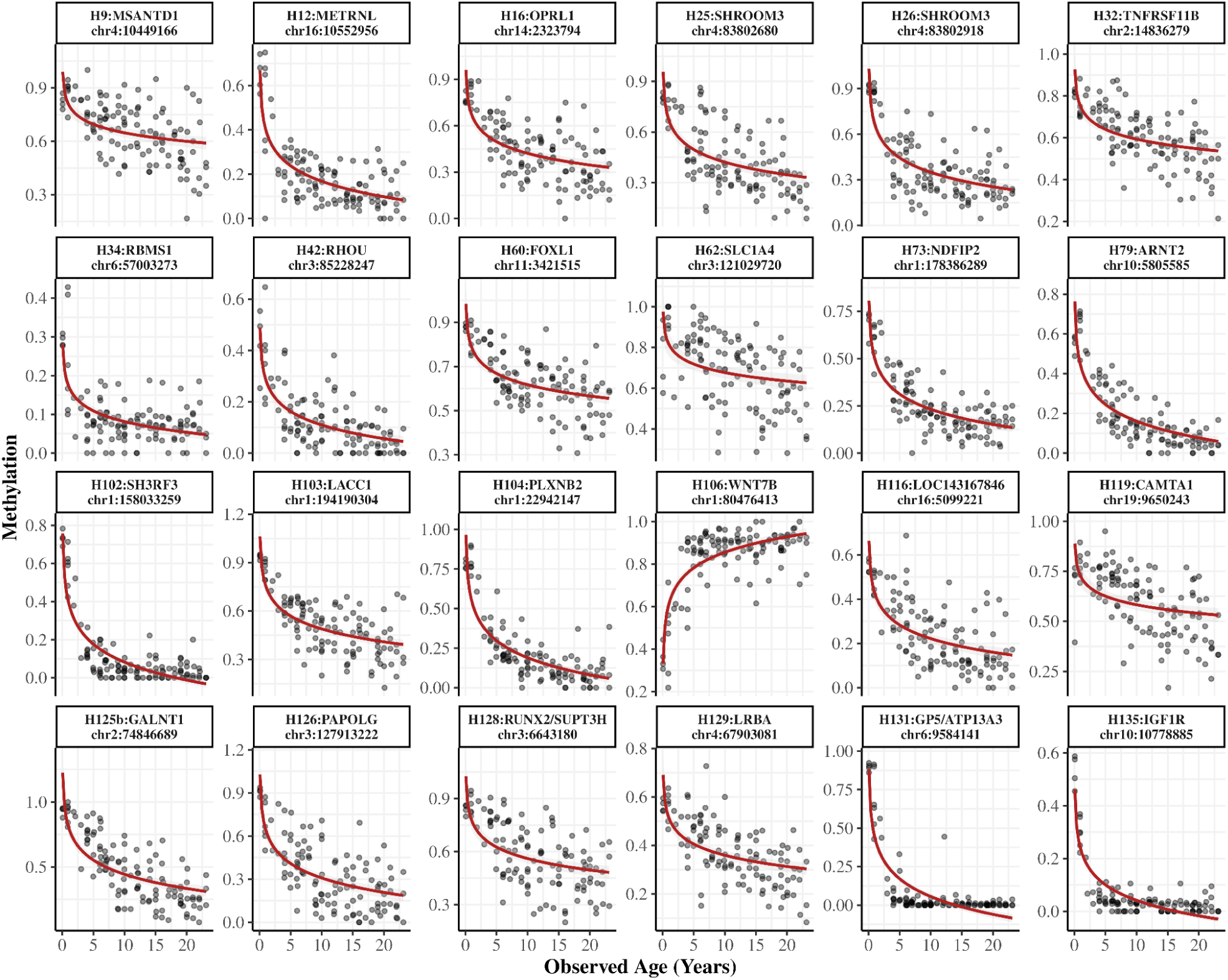
Age-dependent methylation across BEAC-Seq loci in the king penguin. Panels display the proportion of methylated reads (ranging from 0 to 1, y-axis) against observed age (x-axis) for the CpG with the highest adjusted R^2^ with log(Age) per amplicon. Annotations include primer number, gene context, and genomic position, and the y-axis scales vary across panels. A logarithmic smoothing line is applied to each panel to visualize the characteristic flattening of methylation change in older individuals. The plots include methylation data from 119 King penguins (78 EM-Seq samples used for clock training and 41 BEAC-Seq samples used for testing).

**Table 1.** PCR reaction protocols used in primer screening and BEAC-Seq multiplex-PCR. Enzyme MM (Master Mix) corresponds to Invitrogen Platinum SuperFi U Multiplex Master Mix or Qiagen Multiplex PCR Master Mix (top row). MQ refers to Milli-Q water provided in the corresponding kits.

| Component | SuperFi U (Invitrogen) |  | Multiplex PCR (Qiagen) |  |
| --- | --- | --- | --- | --- |
|  | Volume in singleplex-screen (µL) | Volume in BEAC-Seq multiplex (µL) | Volume in singleplex-screen (µL) | Volume in BEAC-Seq multiplex (µL) |
| Enzyme MM | 3.125 | 3.125 | 6.25 | 6.25 |
| Primer F (10uM) | 1.88 (final 1.5 µM) | 1.5 (final 0.06 µM) | 1.88 (1.5 µM) | 1.5 (final 0.06 µM) |
| Primer R (10uM) | 1.88 (final 1.5 µM) | 1.5 (final 0.06 µM) | 1.88 (1.5 µM) | 1.5 (final 0.06 µM) |
| BS-DNA | 1 | 1 | 1 | 1 |
| MQ | 4.62 | 5.38 | 1.49 | 2.25 |
| **Total** | 12.5 | 12.5 | 12.5 | 12.5 |

**Table 2.** PCR cycling conditions used in the BEAC-Seq.

| PCR cycles | Step | SuperFi U (Invitrogen) |  | Multiplex PCR (Qiagen) |  |
| --- | --- | --- | --- | --- | --- |
|  |  | Temperature °C | Time | Temperature °C | Time |
| 1 | Initial Denaturation | 98 | 1:00 | 95 | 15:00 |
| 35 | Denaturation | 98 | 0:15 | 94 | 0:30 |
|  | Annealing | 56 | 5:00 | 56 | 3:00 |
|  | Extension | 72 | 0:30 | 72 | 0:30 |
| 1 | Final extension | 72 | 5:00 | 72 | 10:00 |
| ∞ | Hold | 4 | ∞ | 4 | ∞ |

### An epigenetic clock based on 24 amplicons accurately predicts age in king penguins

Trained using elastic net regression against log-transformed age (see Methods), the BEAC-Model demonstrated high predictive accuracy in the focal population, wild king penguins, both in the training (R^2^=0.88) and the testing (R^2^=0.79) data (Figure 3C-D). To minimize biases, model training was restricted to wild individuals and to one sample per bird^41^. After the removal of 30 zoo-housed and 10 longitudinal replicate samples, and inclusion of 21 additional individuals (see Methods), the final training dataset comprised EM-Seq methylation profiles from 78 wild King penguins (ages 0.88-23.12 years; Figure 3A, Supplementary Table 1). The model was subsequently tested in an independent set of methylation profiles of 41 additional wild king penguins (ages 0.07–22.00 years; Figure 3B) generated via the BEAC-Seq. The mean absolute error (MAE) was 1.71 years for the training data and 2.3 years for the testing data. Prediction error increased with chronological age, with estimates for older individuals showing greater absolute deviations than those for younger birds (Figure 3). This pattern is consistent with the predominantly logarithmic age-methylation relationships observed at many CpG-sites (Figure 2), where methylation changes are most pronounced early in life and become progressively less informative at older ages, while in general epigenetic noise is known to increase with aging, possibly increasing the prediction error in older birds^47^.

**Figure 3.**
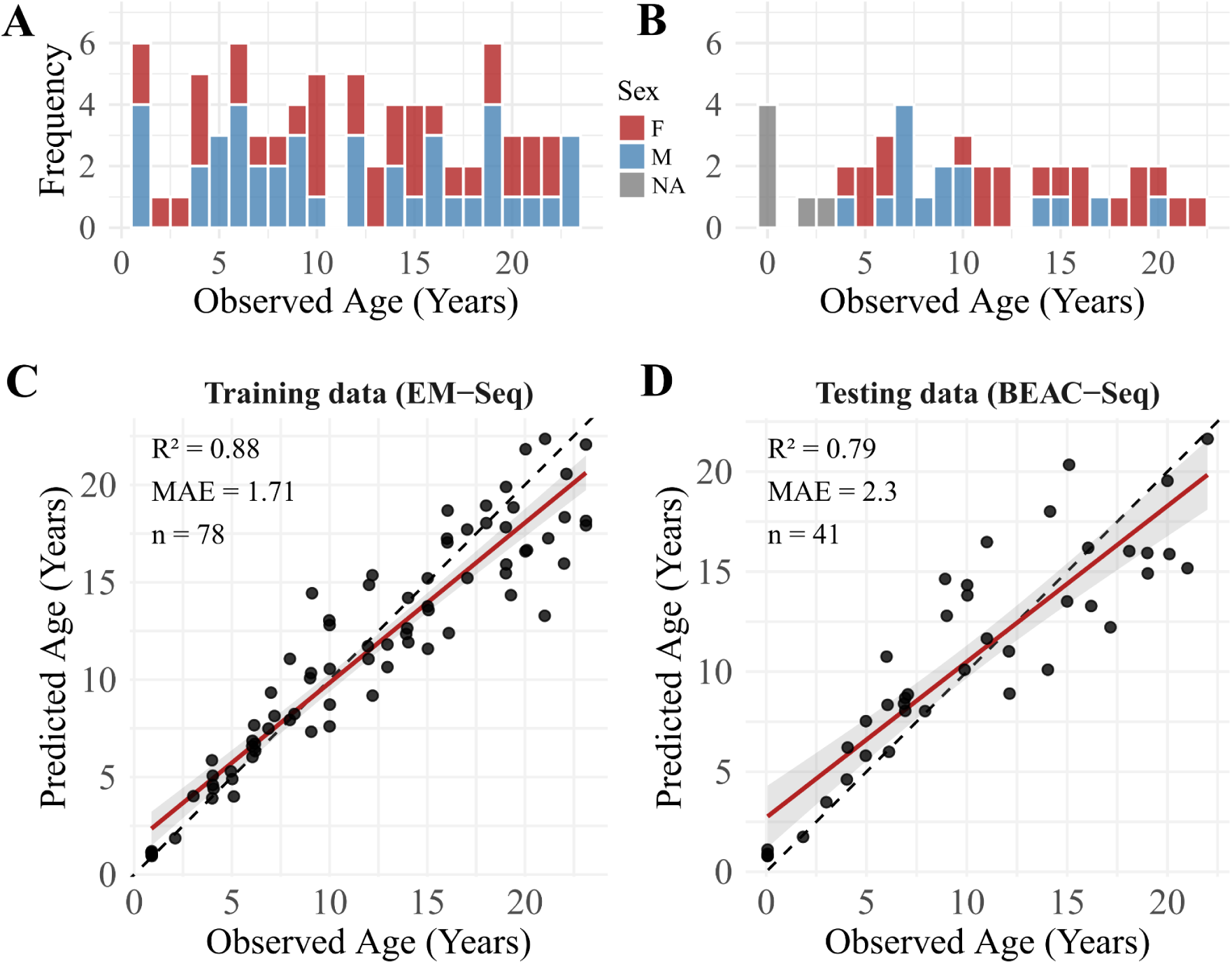
Epigenetic clock performance and validation in king penguins. Distribution of observed age and sex (F = female, M = male, NA = unknown) of the king penguins in the primary training (**A**, n = 78) and testing data (**B,** n = 41). Observed vs. predicted age for the training data **(C)** and testing data **(D)**. In panels **C–D**, dashed lines represent the 1:1 identity line, while the red line indicates the linear regression fit flanked by its 95% confidence interval in grey. Model mean absolute error (MAE), R^2^ and sample size (n) are given for each plot.

The BEAC-Model was trained using elastic net regression with internal cross-validation to optimize the regularization parameters and select the best predictors out of all the available 82 CpG-sites (Methods). This approach selected 33 CpG-sites from 18 different primer pairs as the predictors (Supplementary Figure 1). We also tested how a model trained with generalized additive models^48^ (GAMs) would perform to capture non-linearities between methylation and log-transformed age (Figure 2), using the same training and testing sets as for the elastic net approach. Compared to a GAM fit, the BEAC-Model showed a marginally lower MAE (BEAC MAE=2.3 years, vs GAM MAE=2.39 years), and explained a higher proportion of total variance in the testing set (R^2^ = 0.79 for the BEAC-Model, vs R^2^ = 0.77 for GAM-based clock). Given its edge in both predictive accuracy and explained variance, even if marginal, we ultimately favored the elastic net model for our final framework. Comparison of the BEAC predictors with the GAM approach revealed high consistency: 12 of the 16 influential primer pairs identified by the GAM (effective degrees of freedom >0.01) were also retained in the primary elastic net BEAC-Model (Supplementary Figure 1-2).

### Technical repeatability, longitudinal consistency, and cross-platform bias

The BEAC showed high technical repeatability across different laboratory batches. To evaluate this, ten individual king penguin samples were processed in independent batches (Batch 1 and Batch 2) for bisulfite conversion and Polymerase Chain Reaction (PCR). The resulting age predictions showed remarkable consistency between Batch 1 and Batch 2, characterized by near-perfect alignment (ICC = 0.999; Lin’s CCC = 0.999, Figure 4A). The absolute difference in predicted age between technical replicates ranged from 0.014 to 0.43 years, with a Mean Absolute Difference (MAD) of 0.17 years (Figure 4B). These metrics indicate high absolute precision and negligible laboratory batch-effect noise.

**Figure 4.**
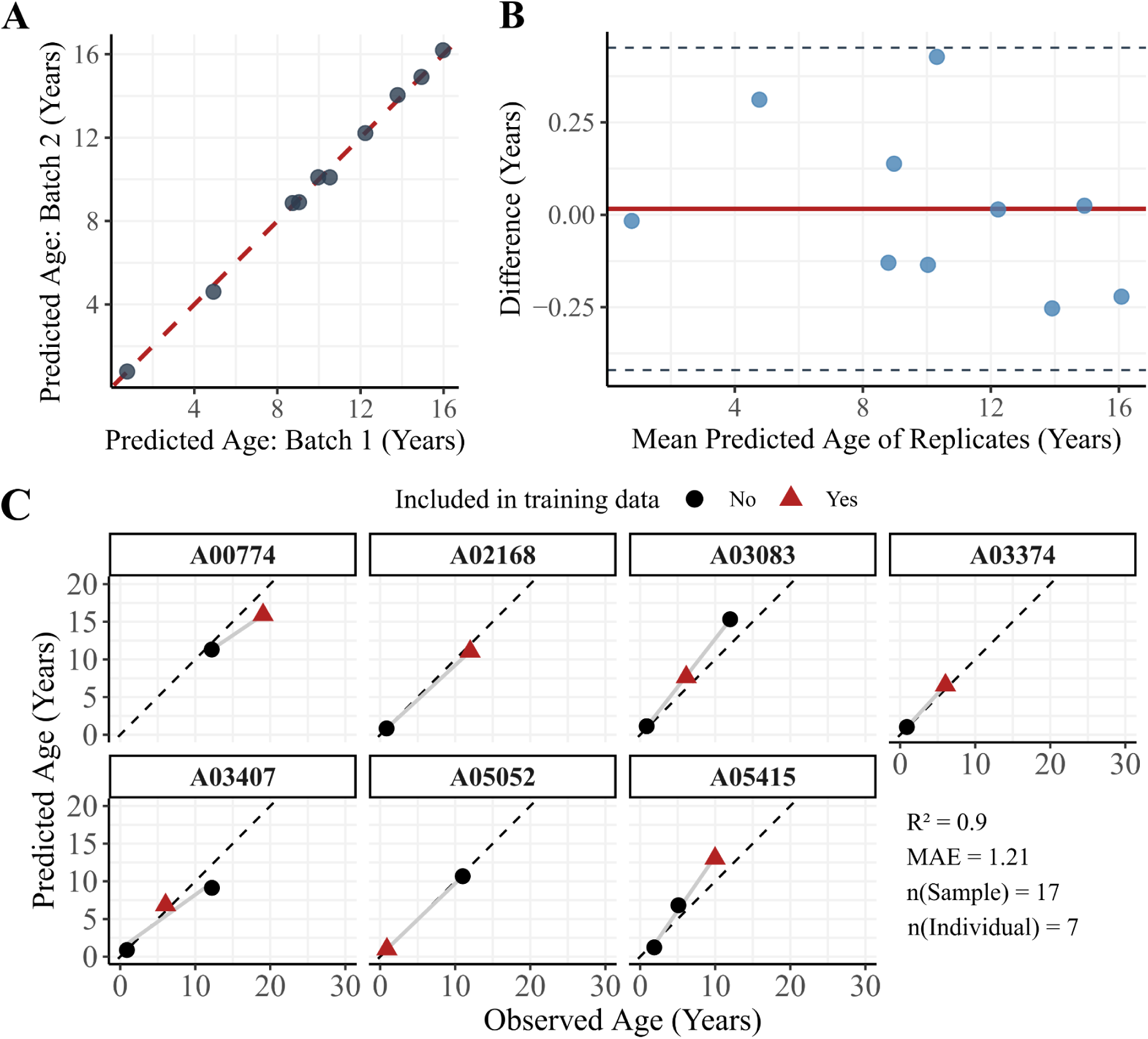
Repeatability evaluation of the BEAC. **(A)** Identity alignment plot comparing predicted epigenetic ages between independent laboratory runs (Batch 1 vs. Batch 2) for ten individual king penguin samples. The red dashed line denotes 1:1 agreement. **(B)** Bland-Altman plot illustrating the distribution of age differences against the mean predicted age of the technical replicates. The solid red line indicates the mean bias (mean difference = 0.05 years), and the dashed red lines represent the 95% limits of agreement (±1.96 SD). **(C)** Longitudinal age predictions for seven individuals across 17 sampling points. Red triangles denote samples utilized in the training data, and dashed lines represent the 1:1 identity line.

To test for model consistency across the life of individual birds, we used repeated longitudinal sampling for seven wild king penguins. For each of the seven individuals, one randomly selected sample had already been included in the initial model training (indicated by red triangles in Figure 4C). Each individual had two to three samples, with ages ranging from 0.9 to 19 years (Supplementary Table 1). Longitudinal EM-Seq samples from the same individuals showed a consistent ageing trajectory: the BEAC-Model achieved a high R^2^ (0.9) and low MAE (1.21 years; Supplementary Table 1, Figure 4C). This demonstrates the model’s ability to reliably track individual biological ageing over time, even across a wide span of years.

We then tested whether training on EM-Seq data introduced a bias in our algorithm, and found that our training/test split did not differ from any random split of our data: across 100 random draws of pooled EM-Seq and BEAC-Seq (pooled n = 119) replicating the original split proportions (78 training, 41 testing), the test data R^2^ and MAE were comparable to our initial results (BEAC: R^2^ = 0.79, MAE = 2.3 years; GAM: R^2^ = 0.77, MAE = 2.39 years). For elastic net, across all 100 random draws, the median R^2^ was 0.74, with 95% coverage interval (CI) of 0.55-0.86 and MAE was 2.46 (95%CI=1.79-3.42), while for GAM models the median R^2^ was 0.71 (95%CI=0.50-0.85) and MAE was 2.82 (95%CI=2.13-3.74; Supplementary Figure 3). This suggests that the BEAC-Model is not overfitting to EM-Seq-specific artifacts (a common form of data leakage in machine learning^49^) but is instead capturing robust biological ageing features that generalizes across platforms.

Median model performance (measured as MAE and R^2^) also remained relatively stable when the training and testing sets were downsampled to 40% of the initial sample, and the model was retrained using elastic net (Supplementary Figure 3A-B). GAMs, however, were much more sensitive to sample size: trained clock models showed significant performance degradation when trained with reduced sample sizes (Supplementary Figure 3C-D). This downsampling analysis was performed by reducing the total pool to 80% (n = 95), 60% (n = 71), and 40% (n = 48) of the original size (n = 119). Crucially, for both models, the spread of the performance metrics narrowed substantially as sample size increased. This convergence indicates that the models become increasingly robust with larger datasets and highlights the high sensitivity of both approaches to training/testing set selection when sample sizes are small.

### Preservation of the age distribution characteristics

To evaluate whether the BEAC-Model accurately captured population-level demographic structure, we compared the distribution of ages predicted by the model with the distribution of known chronological ages in the same individuals. The distributions of observed and predicted ages were statistically indistinguishable based on Kolmogorov-Smirnov (KS) tests for the full dataset (KS D = 0.134, p = 0.232), as well as when the training (KS D = 0.167, p = 0.227; Supplementary Figure 4A) and testing datasets (KS D = 0.171, p = 0.589; Supplementary Figure 4B) were analyzed separately. These results indicate that the model not only predicts individual ages with high accuracy but also preserves the overall age structure of the sampled population, supporting its use in demographic inference and population management applications.

### Resilience to missing data

We demonstrated that our BEAC-Model was resilient to missing BEAC-Seq amplicons, an essential characteristic for real-world experimental conditions where some primer pairs might fail amplification. To evaluate the model’s resilience to data loss, we simulated the absence of amplicons from the BEAC-Seq primer pairs using two distinct sensitivity analyses. First, we tested the original BEAC-Model against single and combinatorial deletions of up to nine out of the 24 amplicons of the BEAC-Seq primer pairs from the testing data (n = 41, BEAC-Seq), imputing missing values via a training-data-based K-nearest neighbors (KNN) panel. Due to the high number of potential combinations (up to 1,307,504 for nine primer pairs), we randomly sampled 1,000 scenarios for combinations of three or more missing amplicons. Single-amplicon deletions demonstrated high stability, with a median R^2^ of 0.79 (95%CI = 0.77-0.80) and MAE of 2.31 years (95%CI = 2.29-2.45, Supplementary Figure 5). Even with nine amplicons missing from the testing data, the model maintained robust performance with a median R^2^ of 0.77 (95%CI = 0.72-0.80) and MAE of 2.54 years (95%CI = 2.32-3.11, Supplementary Figure 5). We performed a targeted “worst-case” simulation by cumulatively removing the nine most influential amplicons, ranked by their total absolute CpG coefficients in the BEAC-Model. Despite losing the most heavily weighted age-associated signals, the BEAC-Model maintained moderate predictive power, achieving an R^2^ of 0.69 and a MAE of 2.92 years even when all nine primary drivers were excluded (Supplementary Figure 6).

**Figure 5.**
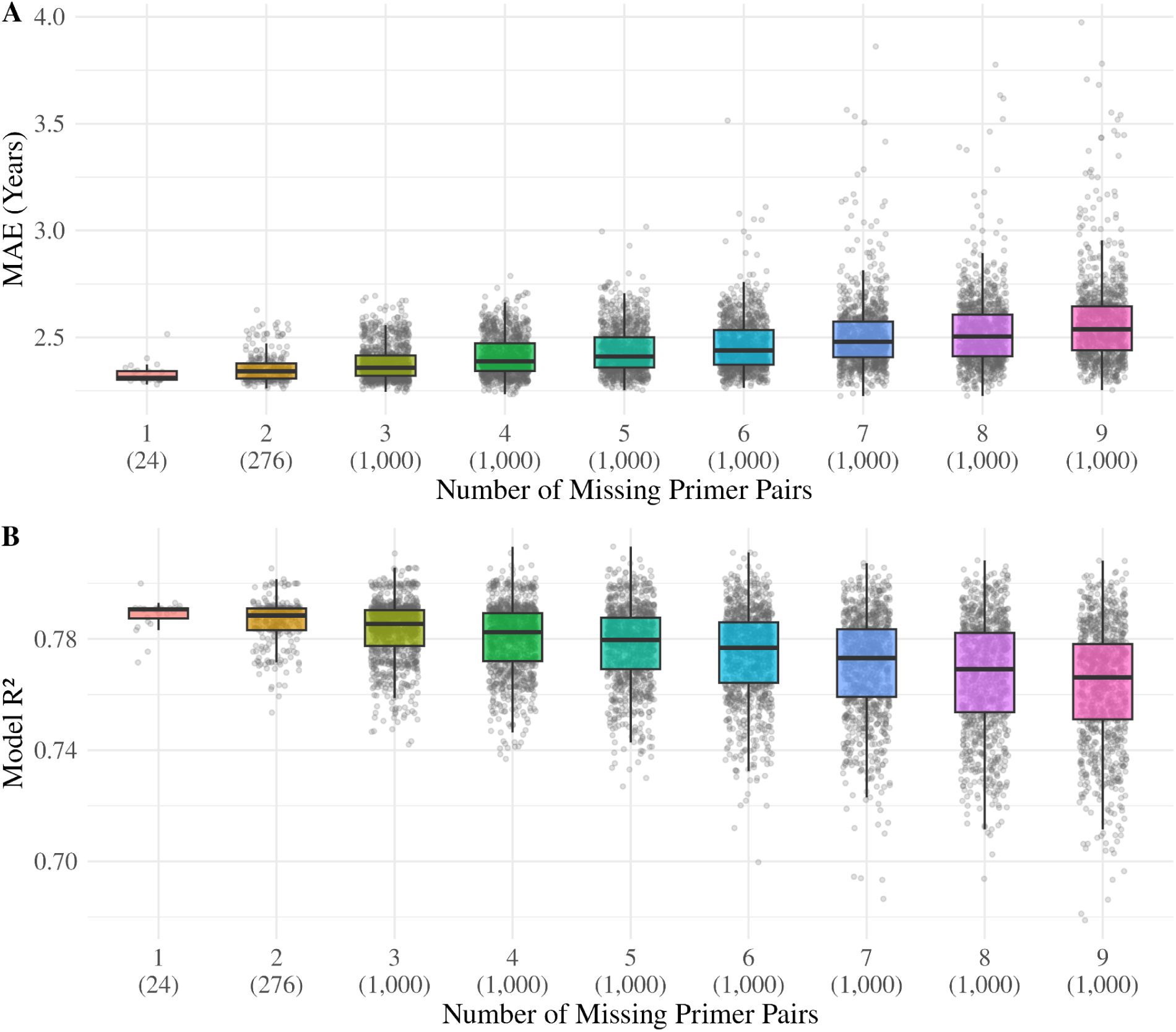
Resilience analysis of the BEAC-Seq. Evaluation of missingness of 1-9 primer pairs of the BEAC-Seq across all possible missingness combinations (1-2 primer pairs missing) and a random selection of 1000 missingness combinations (3-10). The model was iteratively retrained within an elastic net framework using only the CpG-sites of the remaining primer pairs. **(A)** Mean absolute error (MAE, years) and **(B)** R^2^ relative to the number of excluded primer pairs. Values in parentheses indicate the total number of different tested combinations for each count of missing primer pairs.

**Figure 6.**
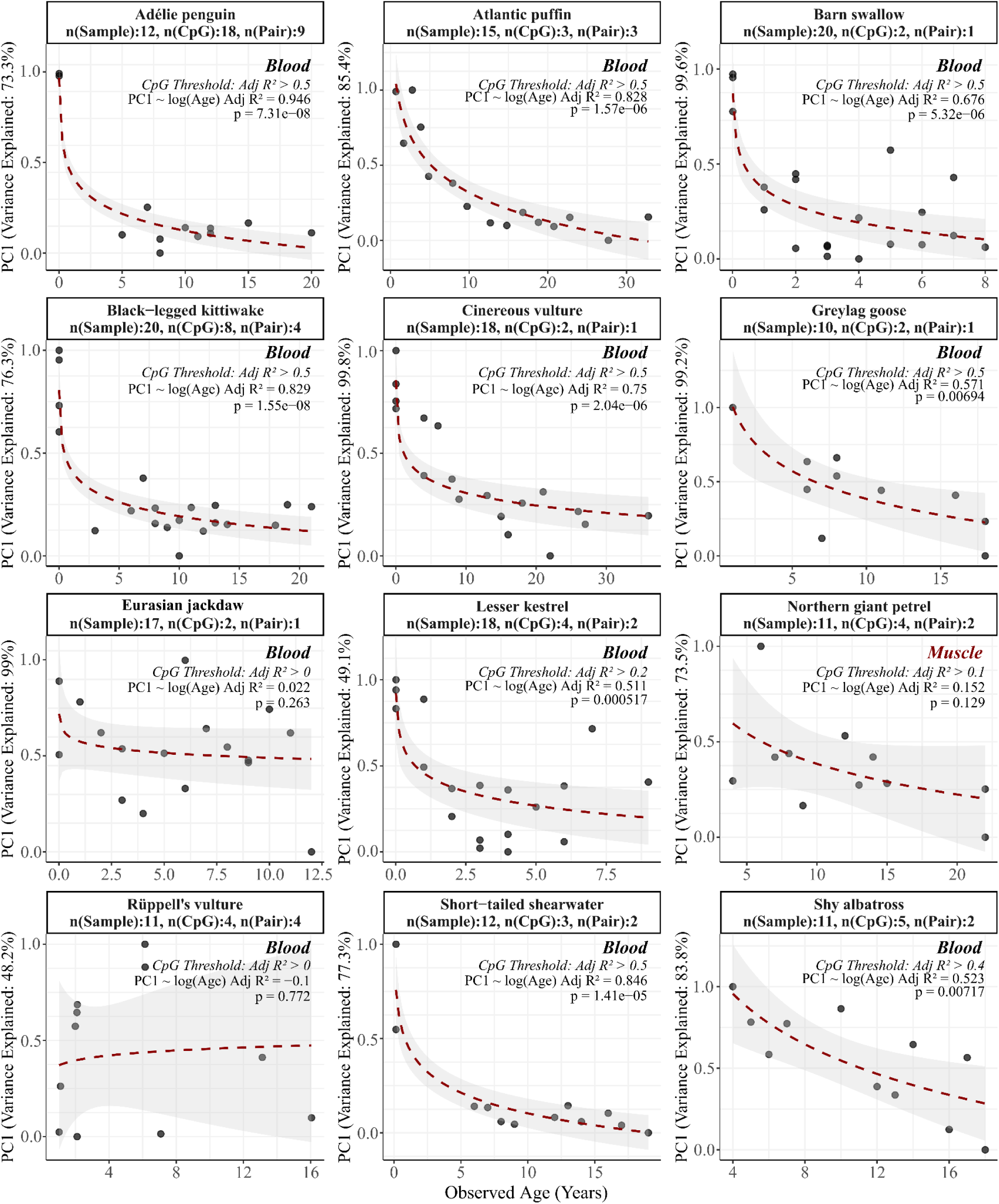
The first methylation principal component by age per species. For each species, input CpG-sites were selected using a progressive adjusted R^2^ threshold (R^2^ = 0.5 down to 0.0, stepping by 0.1). Red dashed lines represent a logarithmic regression fit. Panel margins detail species-specific sample sizes (*n*), filtered CpG and primer counts, baseline selection thresholds, total variance explained by PC1, and adjusted R^2^ values alongside corresponding p-values for the logarithmic age fit for PC1∼log(Age).

Second, we evaluated a retraining strategy across all 24 amplicons from the full panel of primer pairs originally available for model training. For each scenario, ranging from single to 9-amplicon deletions, we retrained the clock using a correspondingly masked training dataset (n = 78, EM-Seq). These retrained models echoed the R^2^ of the simple imputation strategy, maintaining a median R^2^ of 0.77 (95%CI=0.70-0.80), but resulted in consistently lower MAE, with a median of 2.46 years (95%CI=2.24-2.86) even at the maximum deletion level of nine amplicons (Figure 5). These results suggest that applying the BEAC-Seq panel of 24 primer pairs will be robust to experimental data loss. The model can be retrained for the specific use case using our provided retraining script or the imputation panel (n = 119). As our panel derives solely from king penguins, imputation is likely to be less reliable in other bird species, as age-associated methylation signal may not be conserved; cross-species application may therefore benefit from retraining rather than imputation.

### BEAC-Seq detects ageing signals in other bird species

BEAC-Seq primers captured age-related methylation across blood samples of nine bird species (with 11 to 20 samples per species). We designed a subset of the BEAC-Seq primers that can be used or modified to account for DNA sequence polymorphisms in other birds (Supplementary Table 3), and tested them using blood and muscle samples of 15 different species. Overall, primer performance and the relationship between methylation and age varied greatly by species (Supplementary Figures 7–19); while several species yielded an age-predictive signal from at least one tested primer pair, both the direction of methylation change and the proportion of successful primers were highly species-specific. For each species, we selected the CpG-sites with the most age-related methylation signal by adjusted R^2^ (R^2^), subsequently performed a principal component analysis of methylation across the selected CpG-sites, and modeled the first component as a function of observed log(Age) (Figure 6). The variance in the methylation data explained by the first principal component ranged from 48.2% (Rüppell’s vulture, *Gyps rueppelli*) to 99.8% (cinereous vulture, *Aegypius monachus*). We found the best age-predictive capacity in the adélie penguin (*Pygoscelis adeliae*): of the 18 primer pairs tested, 16 successfully detected CpG methylation across 65 CpG-sites, with nine of them (amplifying 18 CpG-sites) having an R^2^ over 0.5 with methylation and age or log(Age) linear models (Figure 6, Supplementary Figures 7-8). The relationship between the first principal component from these 18 CpG-sites from nine primers and log(Age) had an adjusted R^2^ of 0.95 (Figure 6).

In addition to the adélie penguin, promising age-predictive capacity was also observed in eight other species’ blood samples. Individual primers reached an R^2^ >0.5 for the atlantic puffin (*Fratercula arctica*), barn swallow (*Hirundo rustica*), black-legged kittiwake (*Rissa tridactyla*), cinereous vulture, greylag goose (*Anser anser*) and short-tailed shearwater (*Ardenna tenuirostris*, Supplementary Figure 7), with their respective PC1∼log(Age) models yielding R^2^ values between 0.57 (greylag goose) and 0.85 (short-tailed shearwater, Figure 6). For species with weaker individual primer performance, the multi-CpG PC1 approach still provided predictive power: the shy albatross (*Thalassarche cauta*, two primers with R^2^ >0.4) and lesser kestrel (*Falco naumanni,* two primers with R^2^ >0.2) achieved PC1∼log(Age) R^2^ values of 0.52 and 0.51, respectively (Figure 6).

In contrast, the remaining three species showed minimal age-association. The Northern giant petrel (*Macronectes halli*) samples, derived from muscle and not from blood, yielded only two primer pairs including four CpG-sites with R^2^_Adj_ > 0.1 (Supplementary Figure 16) and a non-significant PC1 relationship (Figure 6), a predictable outcome given that the primers were optimized for blood samples. Furthermore, no age correlation was detected across the 11 Rüppell’s vulture samples, a notable contrast to the clear age-related signal found in the closely related cinereous vulture (Supplementary Figure 17 vs 12). Finally, the Eurasian jackdaw (*Coloeus monedula*) showed no significant relationship with age at either the individual CpG site or PC1 level (Supplementary Figure 13, Figure 6).

### BEAC-Seq primer function in additional species

In addition to the primary species used for age-relationship testing, we evaluated primer functionality across other avian species where limited sample sizes precluded age-association analyses. In two Emperor penguins (*Aptenodytes forsteri*, aged 0 > years), the sister species of the king penguin, 22 primer pairs successfully amplified a methylation signal, yielding 86 detectable CpG-sites. Similarly, in two common ravens (*Corvus corax*, aged 5.5 and 8.5 years), three primer pairs (H128, H79, and H73) successfully cross-amplified, capturing five CpG-sites.

## Discussion

This study introduces the Bird Epigenetic Ageing Clock (BEAC), a bisulfite amplicon sequencing protocol and predictive model that offers a scalable, robust, and minimally invasive framework for age estimation in king penguins. Developed from enzymatic methylation sequencing data, the BEAC condenses high-resolution, genome-wide methylation signals into a panel of 24 primer pairs. The panel thus bridges the gap between high-throughput genomic methods and the logistical constraints of field ecology and conservation. To our knowledge, this study introduces the first truly accessible epigenetic clock for birds, as pioneering avian epigenetic clocks so far relied on resource-intensive methodologies DREAM^28^, RRBS^24^, EM-Seq^29^, and WGBS^33^. A subset of the BEAC-Seq primers captures age-related signal across multiple bird species, demonstrating the potential for a single set of loci to serve as a near-universal tool for multi-species age estimation in birds, a result that parallels what has been observed for mammals^14^.

### Age prediction accuracy

The BEAC framework demonstrates enhanced predictive strength and demographic applicability when compared to existing avian epigenetic clocks. For the king penguin, BEAC achieved a robust training R^2^ of 0.88 and a testing R^2^ of 0.79, with an independent testing MAE of 2.3 years, which is equivalent to 7.7% and 5.3% of the species’ maximum observed lifespan in the wild (30 years) and zoos (43 years), respectively. In comparison, earlier frameworks tracking wild adult birds showed lower correlations, though they laid crucial groundwork for the field. This includes the Short-tailed shearwater clock^28^, which was the first epigenetic clock developed for birds, achieving a testing R^2^ of 0.40 across a 21-year lifespan, as well as the subsequent Great tit clock^24^ (0.13–6.03 years) with an LOOCV R^2^ of 0.71. To our knowledge the only two remaining avian clocks - developed for broiler chickens^33^ and Chestnut-crowned babbler nestlings^39^ - report tight errors on the scale of days but only capture narrow developmental windows (under 35 days). Because these early-life clocks cannot track long-term adult ageing, they hold less utility for population demographic modeling. By bridging this gap, BEAC builds upon early avian ageing frameworks to provide the long-term tracking accuracy required for wild populations across decades.

Our development of the BEAC framework complements and expands upon recent breakthroughs in large-scale comparative epigenetics, most notably the pan-mammalian clock developed by Lu et al. (2023)^14^. By utilizing the mammalian methylation array^18^ targeting 36,000 CpG-sites across 185 species, Lu et al. achieved a remarkable median cross-validation correlation (r=0.925; R^2^=0.855). While their model represents a monumental leap in animal ageing studies, it highlights a taxonomic gap: no equivalent universal hybridization chip or clock exists for non-mammalian vertebrates. Our PCR-based, bisulfite amplicon sequencing protocol BEAC-Seq is a step towards closing this gap, capturing age-related signals across multiple bird species, at a fraction of the cost of genome-wide sequencing. Until a dedicated “pan-avian chip” is engineered, the BEAC framework stands as the first accessible, scalable tool, and a pioneering step towards a universal multi-species ageing model for birds.

### Technical robustness

The BEAC-Model demonstrates strong technical robustness and generalizability, supported by the near-perfect agreement between laboratory batches (ICC and Lin’s CCC both 0.999) and the very small mean absolute difference (0.17 years). This exceptionally low batch-to-batch variability answers to recent calls for higher-depth, base-resolution sequencing to overcome technical noise, batch effects, and low reproducibility observed for existing clocks^50^, especially for non-human clocks trained with RRBS data^51,52^. The low reproducibility of clocks trained with RRBS data likely results from inconsistent locus coverage and highly variable sequencing depth across libraries^52^, while CpG-site dropout is also possible in PCR-based bisulfite amplicon sequencing^53^. Sensitivity analyses simulating missing BEAC-Seq amplicons revealed that model performance remains largely stable even under substantial data loss. Notably, even in a worst-case scenario — where the nine most informative primers were removed — the model retained moderate predictive power, indicating that age-related information is distributed across multiple loci rather than concentrated in a few dominant features. Biologically, this supports the view that epigenetic ageing is an interconnected process, characterized not only by targeted alterations at individual CpG sites but also by systemic shifts like global hypomethylation^51,54^. Furthermore, retraining the model on reduced primer sets produced comparable R² and slightly improved MAE relative to imputation approaches, suggesting that adaptive retraining may be preferable when systematic data loss occurs.

Importantly, the model does not appear to overfit platform-specific features, as demonstrated by consistent performance across 100 randomized train-test splits and comparable results between BEAC and GAM approaches. Preventing data leakage, where a model inadvertently accesses information during training that artificially inflates its performance, is a widespread challenge in ecological and biological machine learning that often compromises reproducibility^49^. By strictly isolating our data splits and testing that it is not associated with bias, the BEAC-Model ensures true generalizability rather than leaked training signals. Overall, these results support the robustness and flexibility of the BEAC-Model in real-world experimental settings, while also emphasizing the value of the full primer panel and the availability of retraining or imputation strategies to mitigate missing data.

In addition to evaluation on technical replicates, longitudinal sampling further suggests that the BEAC captures biologically meaningful ageing trajectories within individuals, as reflected by the high R² (0.9). While longitudinal validation is frequently called for in review articles addressing epigenetic clocks to confirm a clock’s biological validity and technical reliability^2,8,55^, such data remains almost exclusively limited to human cohorts^50,56^, and very rare in animal studies^20^. Unlike cross-sectional designs, longitudinal tracking follows individuals across their lifespan, offering a distinct advantage in distinguishing individual phenotypic ageing trajectories from mere chronological time^8^. By assessing longitudinal ageing trajectories, our study addresses a key limitation in the literature and demonstrates the reliability of the BEAC-Model beyond cross-sectional data.

As we evaluated the effect of training data sample size in our downsampling analysis, we confirm that our elastic net model’s predictive stability improves with larger datasets. Although supervised machine learning, such as elastic net regressions using linear or logarithmic transformations, are the standard approach for constructing epigenetic clocks^2^, DNA methylation has been found to exhibit more nuanced, non-linear relationships with age^9,57^. Therefore, we evaluated whether spline-based GAMs would yield more accurate age predictions. Elastic net models produced more accurate predictions, especially at smaller sample sizes, where they proved far more resilient to data reduction than GAMs. Together, these findings indicate that the BEAC-Model captures robust epigenetic ageing signals that generalize across experimental conditions and analytical frameworks, while also emphasising that adequate sample sizes are critical for unlocking the potential of more complex, non-linear models like GAMs.

### Cross-species applicability

The cross-species applicability of the BEAC primer panel further highlights its potential as a flexible tool for avian epigenetic age estimation. A subset of BEAC primers successfully amplified methylation signals in 14 additional bird species, of which 12 had sufficient known-age sample size for evaluation, while insufficient sample sizes precluded testing in common ravens (n = 2) and emperor penguins (n = 2). Among these 12 species, spanning seven out of roughly 35 neognath orders^58^, nine exhibited clear age-associated methylation patterns, demonstrating that the BEAC targets conserved epigenetic loci with relevance across diverse avian lineages. The remaining three species highlight critical methodological and phylogenetic nuances in epigenetic clock application. First, despite their close phylogenetic relationship, the BEAC identified a clear age signal in the Cinereous vulture but failed to do so in the Rüppell’s vulture. This discrepancy may stem from a skewed age distribution; our Rüppell’s vulture samples were aged 1–16 years (with over half aged 1–2 years), and thus did not cover the majority of the species’ 40–50 year lifespan^59^. Second, the non-significant relationship between methylation and age in the Northern giant petrel likely reflects tissue specificity: these samples were derived from muscle tissue, whereas the clock was optimized for blood samples, aligning with the tendency of epigenetic clocks to underperform outside their training tissue^60^. Finally, the weak age signal in Eurasian jackdaws, combined with the fact that both passerine species (Eurasian jackdaw and barn swallow) amplified only a single primer pair across all samples, suggests that the current BEAC panel is not reliably age-informative for passerines. While this single amplicon was strongly age-related in the barn swallow, it lacked informative value in the Eurasian jackdaw (Figure 6) —a divergence that may reflect shifted loci or technical limitations, given the Eurasian jackdaw’s considerably lower DNA yield in our study. Clearly, further work is needed to extend the BEAC framework to passerines.

### Limitations

Several limitations should be considered when interpreting the performance and broader applicability of the BEAC-Model. First, our initial CpG-site selection was limited to king penguins, introducing an initial species-specific bias. The array-based pan-mammalian clock used at maximum 174 species in elastic net marker discovery, and was thus able to capture highly conserved ageing signals^14^. While no such pan-species chip exists for birds, and we identified candidate age-related CpG-sites solely from genome-wide methylation data of the king penguin, other avian species may harbor additional, more informative age-associated loci not captured by our current primer set. Furthermore, the reliance on a relatively small subset of shared loci across species may constrain maximal predictive accuracy outside the focal taxa. Uncertainty in age estimates, particularly in wild populations where exact hatch dates are often unknown, also introduces unavoidable noise that may affect model training and evaluation. For example, while the ages of the king penguin samples were known to within ∼1 month, human and traditional model organism clocks are typically trained on data with ∼1 day age accuracy. Furthermore, the current model is derived from a single population, which may limit its generalizability across geographically or environmentally distinct groups. For example, sampling population has been found to influence DNA methylation levels in humans^61^, barn swallows^62^ and Atlantic cods (*Gadus morhua*)^25^. Although epigenetic clocks are largely robust to population differences^25,63^, and king penguins show extremely low genetic diversity across populations^64^, minor variations may still affect prediction accuracy across different groups^63^. On the other hand, our 15 Atlantic puffin samples were obtained from five different zoo locations, yet we still detected a strong age-predictive signal for that species, highlighting the potential robustness of these epigenetic markers despite distinct environments. Environmental and physiological factors, such as seasonal effects (e.g., metabolic shifts^41^), could also influence methylation patterns and potentially confound age predictions. While the use of a single tissue type simplifies sampling and contributes to model precision, it may limit performance in tissues such as muscle, as was found for Northern giant petrel samples. Finally, the BEAC-Model should be viewed as an iterative framework: continued expansion across species, tissues, and ecological contexts will be essential to refine marker selection, reduce bias, and improve overall performance.

### Future perspectives

Future work should expand the BEAC framework across both biological and practical dimensions. Beyond age estimation, the BEAC approach also holds potential for exploring how environmental stressors, such as nutritional limitations^65^, environmental toxins (e.g., DDT)^66^, climate variability^66^, or viral infections^67,68^, may influence epigenetic ageing patterns in birds, offering a possible, though still tentative, avenue for linking physiological condition with ecological pressures. While it may be premature to suggest that such tools will revolutionize population management, they could provide valuable insights for demographic monitoring — a crucial factor for long-term conservation outcomes^69^. Specifically, age-determination tools can help the field of socio-ecological studies leap beyond broad morphological age classes (e.g., chick vs adult) toward precise age estimates throughout the lifespan, enabling studies on age-dependent behaviors as well as age-selective threats, such as hunting^70^. Traditional avian age determination methods often rely on expensive and labor-intensive recapture or long-term tracking programs, or morphology such as plumage, size, or vocalisations, which are typically limited to coarse life-stage classification, and frequently lose discriminatory power after maturity^71,72^. In contrast, epigenetic clocks provide quantitative age estimates across adulthood and old age from minimally invasive samples, offering substantially finer temporal resolution for demographic and ecological analyses^71^. This capability is particularly critical for long-lived species like seabirds, which face severe population pressures from cumulative stressors^73^, including mass mortality events driven by fisheries bycatch^74^ and the recent emergence of highly pathogenic avian influenza (HPAI) H5^75^. Because the demographic consequences of these impacts depend heavily on the affected age and sex classes, accurate population-level evaluations require precise age estimation^76^. In sum, epigenetic clocks offer a reliable method for determining chronological age, significantly expanding the scope of population monitoring and demographic modeling. Furthermore, the distributional alignment observed in this study demonstrates that this framework can be reliably deployed as a practical tool for downstream demographic forecasting and wildlife management^77^.

In addition to the direct application of the BEAC in birds, this study serves as a practical framework for epigenetic clock development: by combining genome-wide discovery, SNP-informed primer design, and robust statistical modeling, we provide a transferable “blueprint” that can be adapted to other taxa. The BEAC-Model is not only a species-specific tool, but also an example of how epigenetic clocks could be developed for other non-mammalian vertebrates using the principles described here. The BEAC-Seq panel may directly be applied to other bird species as a low-cost alternative to *de novo* clock development. Because it is a targeted PCR assay, it can be applied without the costly whole-genome sequencing typically required to discover age-associated markers. Our cross-species results show that the primers can recover an age signal in several birds. We recommend that researchers adopt primer sets from phylogenetically related species where possible (Supplementary Table 2-3, Supplementary Data 1), followed by validation against the target species’ reference genome to account for potential sequence variation. For researchers wishing to design custom primers, we recommend aligning our provided FASTA sequences (encompassing the 400bp regions surrounding the 24 target BEAC-CpG-sites in the king penguin reference genome, Supplementary Data 2) against a specific species’ genome to design *de novo* primers around these age-informative loci following the steps described here. Turning this into a working clock will require validation in a larger known-age cohort per species and subsequent recalibration. For empirical validation, our downsampling analysis indicates that singleplex amplification on bisulfite-converted blood-based DNA using a minimum of ∼48 known-age individuals maintains a reliable median R^2^ and MAE (Supplementary Figure 3), providing a sufficient foundation for recalibrating species-specific clocks via elastic net regression using our provided scripts (Supplementary Data 3). For optimal clock training, the sample set should represent a uniform age distribution across the species lifespan. Depending on phylogenetic proximity and tissue type, both direct application of the BEAC and BEAC-CpG-site based primer design has potential to offer a rapid, cost-effective method to epigenetic age estimation across non-model avian species.

## Conclusions

The BEAC-Model provides a robust, flexible, and empirically validated framework for epigenetic age estimation that balances methodological rigor with practical applicability. By demonstrating high technical repeatability, resilience to missing data, cross-species transferability, and relatively low error relative to lifespan, this approach moves beyond proof-of-concept toward real-world usability. Although limitations remain, particularly regarding taxonomic breadth, environmental influences, and population-specific biases, the BEAC framework is inherently iterative and well-suited to refinement as additional data become available. The BEAC framework helps bridge the gap between complex genomic modelling and the practical demands of large-scale ecological and conservation research, providing a tool that is both scientifically rigorous and accessible for further development and broader application.

## Methods

### Software

Specific softwares and packages used in the analysis are detailed in the corresponding method sections. All plots were generated in R with the package ggplot2^78^ and further edited using Inkscape (version 1.4.2)^79^ and BioRender.com.

#### 1. Sample description

King penguin 149 whole blood samples were obtained from known-age king penguins. These king penguins were from a free-ranging population in the Crozet Archipelago (Possession Island), French Southern Territories (n = 119), and European zoos, and housed at Zoo Zürich, Switzerland (n = 10), and Loro Parque, Spain (n = 20). We used 96 samples from 87 King penguins for marker discovery (57 wild, 30 zoo-housed). Of the 87 king penguins used for marker discovery, 57 wild penguins were retained for model training. We additionally generated EM-Seq data for 21 wild individuals for model training, resulting in a training set of 78 samples. For model testing we generated data for 41 additional wild king penguins using BEAC-Seq. Supplementary Table 1 describes samples used in different analysis steps with their corresponding distributions of species, age distribution, sex, origin and rearing environment.

The wild individuals were followed since fledging (1-year-old) as part of a long-term monitoring project^80^, and their sampling was approved by the French ethics committee (latest: APAFIS#29338-2020070210516365) and the French Polar Environment Committee. Permits to handle the animals and access the breeding sites were issued by the “Terres Australes et Antarctiques Françaises” (section Ethical permits). In zoos, samples were collected for routine veterinary examination as part of Zoo Zürich and Loro Parque’s husbandry licences, their reuse was not subject to additional authorisation. Monitoring and sampling processes are further described in Cristofari et al^41^. DNA was extracted from whole peripheral blood samples using a spin-column method (Macherey-Nagel Nucleospin Blood kit), with RNAse A treatment. DNA concentration was measured using a NanoDrop 1000 (Thermo Scientific).

#### 2. Enzymatic sequencing and data processing

To generate the methylome libraries, we utilized the NEB EM-Seq kit for whole-genome enzymatic-conversion sequencing. The full details of sequencing and data processing are described in Cristofari et al^41^. Briefly, libraries were sequenced on an Illumina NovaSeq 6000, and raw reads were quality-trimmed and filtered by length. Processed reads were aligned to the king penguin reference genome using BSBolt^81^ (v1.5.0), followed by deduplication and filtering via SAMtools^82^ (v1.16.1). Finally, average CpG methylation levels were determined by pooling strand-specific calls, and known SNP positions were masked using a genomic database.

#### 3. CpG selection

A total of 96 samples from 87 King penguins were used for clock CpG selection through a three-step filtering process. First, we included only CpG sites with an average sequencing depth across all samples of 20-50 reads per CpG (after pooling strand-specific methylation calls). Second, we excluded individual observations where sequencing depth was below 10 or above 100 for any sample. Third, we retained only CpG-sites where the standard deviation of DNA methylation was above 0.1. This resulted in a final set of 2,429,249 CpG sites. To identify CpG-sites with the strongest relationship to chronological age, we correlated methylation values at each site with age at blood sampling using both Pearson and Spearman correlations. CpG-sites were ranked based on correlation p-values, and the top 100 sites from both correlation methods were selected for further analysis. The overlap between the correlation methods included 82 CpG-sites, resulting in a total of 118 CpG-sites.

#### 4. Primer design

We designed primer sets for the 118 candidate CpG-sites strongly associated with chronological age (Pearson or Spearman). We extracted the DNA sequence from a region 200 bp up- and downstream of each of the candidate CpG-site from the king penguin reference genome. We used BLAST^83,84^ to ensure all CpG-site candidate sequences of 400 bp had only one alignment hit against the reference assembly. PrimerSuite^85^ was used to design bisulfite primers with amplicon lengths of 100-150 bp, a target melting temperature (T_m_) of 56°C, and no CpG-sites within the primer sequence. From the PrimerSuite output, we selected pairs that had no homopolymers (considered at minimum six of the same base, or minimum four of the same base at the 3’ end), while we also aimed to maximize the dimer score. We also aimed at selecting primers with higher GC content (>25%), and a GC-clamp in the 3’ end. We combined the primer candidate information with a SNP panel from 64 genotyped king penguin individuals, and ensured no primers had known SNPs within them. As our aim was to build an epigenetic clock for king penguins that would work well for other bird species, we ranked the candidate primer pairs based on sequence similarity across other bird species using PhyloP^86,87^ scoring. The PhyloP score was calculated based on a reference-free alignment of 363 bird species^46^, to which we added the king penguin genome using Cactus (2.8.1)^88^ and HAL toolkit^89^. For each primer (forward and reverse), we calculated the average PhyloP score across the primer sequence using a neutral model of sequence evolution based on repetitive sequences of the genome, available in Feng et al.^46^, where higher scores indicated higher sequence conservation. Tested primers were selected based on all the above-described parameters: homopolymers, dimer score, GC content, GC-clamp, SNP, and PhyloP score.

Based on PCR performance, we iteratively redesigned a subset of primers to optimize PCR amplification using PrimerSuite as described above. To guide this second iteration, we modeled primer success against several biophysical properties. Shorter primers with higher GC content were strongly associated with improved performance (length: estimate = −0.131, SE = 0.025, p = 3.72*10^−7^; GC: estimate = 0.039, SE = 0.007, p = 1.53*10^−7^; Supplementary Figure 20A–B), a correlation driven by maintaining a stable target T_m_ of ∼56°C (Supplementary Figure 20C). In contrast, predicted T_m_ (estimate = −0.098, SE = 0.30, p = 0.742) and amplicon length (estimate = 0.001, SE = 0.009, p = 0.866, Supplementary Figure 20D) did not significantly impact primer performance, consistent with the narrow T_m_ range imposed during primer design. Both parameters were controlled to ensure downstream multiplex compatibility and uniform sequencing coverage.

To design cross-species primer variants, we extracted alignment files from the above described 364-bird alignment using HAL toolkit^89^. For each tested species, the primers were designed using the closest relatives available in the 364-bird alignment (Supplementary Table 4). For primers that were successful in our initial screenings with low-frequency polymorphisms, we substituted bases to match conserved avian clade patterns. Additionally, we incorporated degenerate bases at positions where polymorphisms resulted in a CpG site within primer binding regions to prevent allelic amplification bias and subsequent skewing of epigenetic data. In total, 139 primer pairs, including these cross-species variants, were designed and tested.

#### 5. Bisulfite-DNA preparation

Bisulfite (BS)-conversion was done using the EZ DNA Methylation-Gold kit (Zymo) using ∼500 ng input DNA, following the manufacturer’s instructions. The BS-converted DNA was stored in −80°C after conversion and the PCR was performed within two weeks of the conversion for all samples.

#### 6. Primer screening

We screened the performance of the primers using bisulfite-converted king penguin DNA. We first screened the performance of the 139 primer pairs in singleplex PCR reactions using bisulfite-converted DNA pooled from ten king penguin individuals. We used either the Invitrogen Platinum SuperFi U Multiplex Master Mix (Fisher Scientific), or Multiplex PCR Master Mix (Qiagen; following the discontinuation of SuperFi U) for the PCR reactions. For both kits, we followed the manufacturer’s instructions with the following exceptions: the total sample volume of the PCR reactions was 12.5μL, and the PCR reaction protocols and cycling conditions were modified as shown in Tables 1 and 2. The individual primer pairs were screened at 1.5μM final molarity. For both enzymes, we tested annealing times of 30 seconds, 1, 3, and 5 minutes, and annealing temperatures of 56°C, 57°C and 58°C. The final primer screening reaction protocols are described for both enzymes in Table 1 (*Volume in singleplex-screen*), while the final PCR cycling programs are described in Table 2. We assessed the amplification of each primer pair using gel electrophoresis (1% agarose gel, 100V) where successful amplification was identified as visual bands at the expected product size. Second, we tested the PCR performance of the primer pairs with successful PCR amplification (89 primer pairs) with TruSeq sequencing adapters (Forward: 5’-ACACTCTTTCCCTACACGACGCTCTTCCGATCT-[Forward primer]-3’; Reverse: 5’- GTGACTGGAGTTCAGACGTGTGCTCTTCCGATCT-[Reverse primer]-3). We used pooled bisulfite-converted DNA from ten king penguin individuals to screen the performance of these primers in singleplex PCR as described above.

#### 7. Multiplex-PCR optimization

We assessed which of the overhang-primers were prone to forming inter-pair heterodimers, and thus interfere with target template amplification in multiplex-PCR. We used PrimerPooler software (v1.89)^90^ to identify optimal subpools of the primer pairs. We divided the 24 primer pairs into two subpools of 12 pairs based on PrimerPooler predictions (Supplementary Table 2). As in the previous primer screening steps, we used 1 μL of bisulfite-DNA per reaction, in a total reaction volume of 12.5 μL. We screened the performance of the multiplex reactions across a range of primer molarity (0.024, 0.032, 0.04, 0.048, 0.055 and 0.06 μL), annealing time (three and five minutes) and annealing temperature (56°C, 57°C and 58°C) gradients. Tables 1 and 2 describe the PCR protocols and cycling conditions in our final BEAC-Seq protocol. After the two-pool multiplex PCR, we pooled the products of the two pools for each sample for sequencing. When extending this protocol to other species, if there were 12 or fewer suitable primers for a specific species’ samples, they were processed in a single multiplex-PCR reaction regardless of within-pool compatibility rather than across multiple pools.

#### 8. Amplicon sequencing

The pooled multiplex products of the BEAC-Seq for each sample were first cleaned using Exonuclease I and FastAP Thermosensitive Alkaline Phosphatase (Thermo Scientific), and subsequently barcoded using in-house TruSeq indexing primers and Phusion Hot Start II DNA Polymerase (Thermo Scientific), and paired-end sequenced on MiSeq and Element Aviti platforms (Illumina). Raw paired-end reads were processed using Trimmomatic^91^ (v0.39) to remove adapter sequences and low-quality bases. Only paired-end reads with a minimum residual length of 36 bp were retained for downstream analyses. Bisulfite-aware alignment was performed using BSBolt^81^ against the king penguin reference (GCA_010087175.1). BAM files were filtered using SAMtools^82^, retaining only primary alignment with MAPQ ≥ 30. Site-specific methylation levels were extracted using BSBolt. We retained only CpG-sites with a total read depth exceeding 30 reads.

#### 9. BEAC-Model development

##### Data preparation

We extracted the same CpG-sites from EM-Seq and BEAC-Seq data using GenomicRanges92 in R (version 4.4.2 used throughout the manuscript)93. We filtered the methylation matrix to exclude CpG-sites with a missingness rate above 15% of the samples in the specific dataset. The resulting filtered matrix was then imputed using the k-nearest neighbors (k-NN) algorithm (k = 10) using the R package impute94 separately for both EM-Seq and BEAC-Seq data. Across all CpG-sites and all individuals, five out of 3,362 observations were imputed for the BEAC-Seq data while ten out of 6,396 observations were imputed for the EM-Seq data. Following the quality control and coordinate synchronization steps described above, we retained a final set of 82 high-quality CpG-sites across 24 primer pairs that were included in both BEAC-Seq and EM-Seq data.

##### BEAC-Model training

The BEAC-Model training set included one sample per individual from 78 King penguins. Although marker discovery used 87 birds (57 wild and 30 zoo-housed), model training was restricted to wild individuals and one sample per bird. An additional 21 wild penguins were sequenced by EM-Seq, yielding the final training set of 78 penguins. The model was trained exclusively on the EM-Seq dataset, utilizing the methylation proportions of the 82 CpG-sites retained in both the EM-Seq and BEAC-Seq data. We used elastic net to regress CpG methylation against log(Age), using glmnet^95^ and glmnetUtils^96^ in R. To optimize the model’s hyperparameters, the mixing parameter (α) and the regularization penalty (λ), we performed a grid search using 10-fold cross-validation on the training set using the function cva.glmnet() from R package glmnetUtils^96^. The optimal α and λ were selected based on the values that minimized the Mean Squared Error (MSE). The final clock was then constructed using these optimized parameters.

##### BEAC-Model testing

To assess the resilience and practical utility of the clock, we performed an external validation on an independent dataset generated via BEAC-Seq. Predictive accuracy was evaluated by comparing the predicted ages (back-transformed from the log-scale) against observed chronological ages. Model performance was quantified using R^2^ and mean absolute error (MAE) to provide a direct interpretation of average age-prediction error in years.

#### 10. Model comparison

##### Generalized additive models

We evaluated the performance of an alternative clock algorithm based on Generalized Additive Models^48^ (GAMs) using mgcv^97,98^ in R. For each amplicon, we first fitted a series of univariate GAM regressions for each single CpG-site against log(Age), and selected each time the single most informative CpG-site based on adjusted R^2^, using EM-Seq data as training data. The GAM-based clock was then fitted on these 24 CpG-sites, with one smooth term per site using Restricted Maximum Likelihood (REML)^99^. As for the elastic net approach, we tested the model on the BEAC-Seq data and evaluated the performance using R^2^ and MAE.

##### Bootstrapping and Iterative Subsampling

We performed a series of downsampling experiments to simulate varying levels of data availability for both the elastic net and GAM versions of the BEAC-Models described above. The original dataset was downsampled to three different proportions: 80% (n = 95), 60% (n = 71), 40% (n = 48) of the original size (n = 119). For each downsampling level, we conducted 100 iterations of a randomized training-validation pipeline. In each iteration, the data were randomly partitioned into a 66% training set and 34% test set to conform to our initial train-test proportions. For elastic net models we re-optimized the hyperparameters (α and λ) via 10-fold cross-validation on the training subset as described above. For GAMs, we refit the splines using REML and automated smoothing parameter selection. For both modeling strategies, within each of the 100 iterations per downsample level, the resulting models were used to predict the age of the held-out test samples, from which we calculated the R^2^ and MAE.

#### 11. Technical resilience

##### Technical repeatability

To assess the technical repeatability and susceptibility to batch effects of the BEAC, we processed ten king penguin samples in independent batches of bisulfite-conversion and multiplex PCR. We evaluated the statistical consistency and replication accuracy between Batch 1 and Batch 2 using three complementary metrics. First, Lin’s Concordance Correlation Coefficient (CCC; epiR^100^ package) was used to assess both directional precision and absolute accuracy relative to a 1:1 identity line. Second, to partition biological variance from technical laboratory noise, we calculated a two-way random effects, single-measure Intraclass Correlation Coefficient (ICC(2,1)) for absolute agreement (irr^101^ package). Finally, the Mean Absolute Difference (MAD) was calculated to define the baseline technical error distribution in absolute biological years.

##### Resilience to missing primers

To evaluate the stability of the epigenetic clock and its practical reliability in the event of technical failures or data loss, we conducted resilience analyses simulating nine levels of missingness (from k = 1 to k = 9 of the 24 BEAC-Seq primer pairs) and tested model recovery by two distinct strategies: data imputation and model retraining.

In the imputation strategy, for each level of missingness, we masked either a combination of k random primer pairs up to 1000 combinations, or the k most influential primer pairs from the testing data. In that case, we quantified the “influence” of each primer pair as the sum of the absolute values of the coefficients of all CpG-sites contained within that pair in the primary BEAC-Model. These missing values were then recovered via k-NN imputation^94^ using the training (EM-Seq) data as a reference. We used the primary EM-Seq-trained BEAC-Model to predict the ages from this recovered testing data, and re-calculated the distribution of R^2^ and MAE across all combinations for each level of missingness to determine how the loss of primary predictive markers influenced the deviation from chronological age.

In the retraining strategy, we repeated the approach above for data deletion, but instead of imputing missing values, we re-trained the model on the fly and tested the performance of the re-trained model. In each combination, the selected primers were masked as missing data in both testing and training data. We then retrained the model using only the remaining primers and used that to predict the ages, and calculated the distribution of R^2^ and MAE for the testing data across all tested combinations for each level of missingness.

#### 12. Extension to additional avian species

To assess the broader applicability of our primer sets, we tested primers adapted for clade-specific polymorphisms (Supplementary Table 3) across blood and muscle samples from 14 avian species other than the king penguin (Supplementary Table 1). The bisulfite conversion and PCR amplification followed the BEAC-Seq protocol, utilizing the primer combinations listed in Supplementary Data 1. All species-specific PCRs were performed in a single pool, with the exception of the adélie penguin, which utilized 16 primers split into two sub-reactions, mirroring the king penguin pooling strategy.

Data processing remained consistent with the king penguin pipeline, with the exception of the alignment and methylation calling stages: reads for each species were aligned to their own reference genome or that of the closest available relative (Supplementary Table 4). To precisely locate target regions in diverse genomes, we employed a custom mapping pipeline using a 400-bp region surrounding each target CpG from the king penguin reference genome as a query sequence. These sequences were filtered to match only the primers utilized for each specific species and then aligned to the target genome using minimap2^102^ (v2.28) with the -a flag to generate high-quality SAM alignments. Alignments were sorted with SAMtools^82^ and converted to BED format using BEDTools^103^, allowing us to define the most likely genomic coordinates for each amplicon in every species. Finally, we used these species-specific genomic coordinates to map the observed methylation calls back to their corresponding primers. In instances where minimap2 failed to predict a plausible genomic location for the 400-bp king penguin query sequence (possibly due to incomplete or highly fragmented reference genomes in the target species) we implemented a fallback alignment strategy. We directly aligned the raw sequencing data against a database of known primer sequences to identify target regions. The process was conducted in R using the package Biostrings^104^ as follows: Overlapping reads from the raw BAM files were clustered to generate a high-confidence consensus sequence for each potential locus using a 50% majority rule. We performed local pairwise alignments between these consensus sequences and a library of forward and reverse-complement primer sequences. A sequence was formally defined as a “successful amplicon” only if both the forward and reverse primers (or their respective complements) were identified within a span of 500bp on the same consensus scaffold.

The final methylation data for each species was filtered to retain only those CpG-sites present in more than three samples. To avoid confounding effects from potential SNPs or technical artifacts, we excluded CpG-sites where more than half of the individuals exhibited extreme methylation values (exactly 0 or 1). The above filtering steps were conducted for all species except for Emperor penguin and Common raven due to low sample size (n = 2).

To assess the maximal cross-species age-predictive capacity of the BEAC-Seq primers given the sample size, an adaptive filtering pipeline was executed across an adjusted R^2^_Adj_ step-down sequence from 0.5 to 0.0 (in 0.1 decrements). For each species, single-site methylation data were evaluated against chronological age using linear (Methylation∼Age) and logarithmic (Methylation∼log(Age)) regressions. To ensure structural viability, CpG loci with more than 15% missing data within a species cohort were removed. Remaining missing values within these validated matrices were resolved via k-Nearest Neighbors (kNN) imputation^94^.

The imputed, centered, and scaled methylation matrices were analyzed using Principal Component Analysis (PCA) to extract the first principal component (PC1). The global age-predictive signal of PC1 was then modeled using ordinary least squares regression against log-transformed age: PC1∼log(Age + 0.01). A baseline offset of 0.01 years was added to prevent errors for samples aged 0.

## Ethical permits

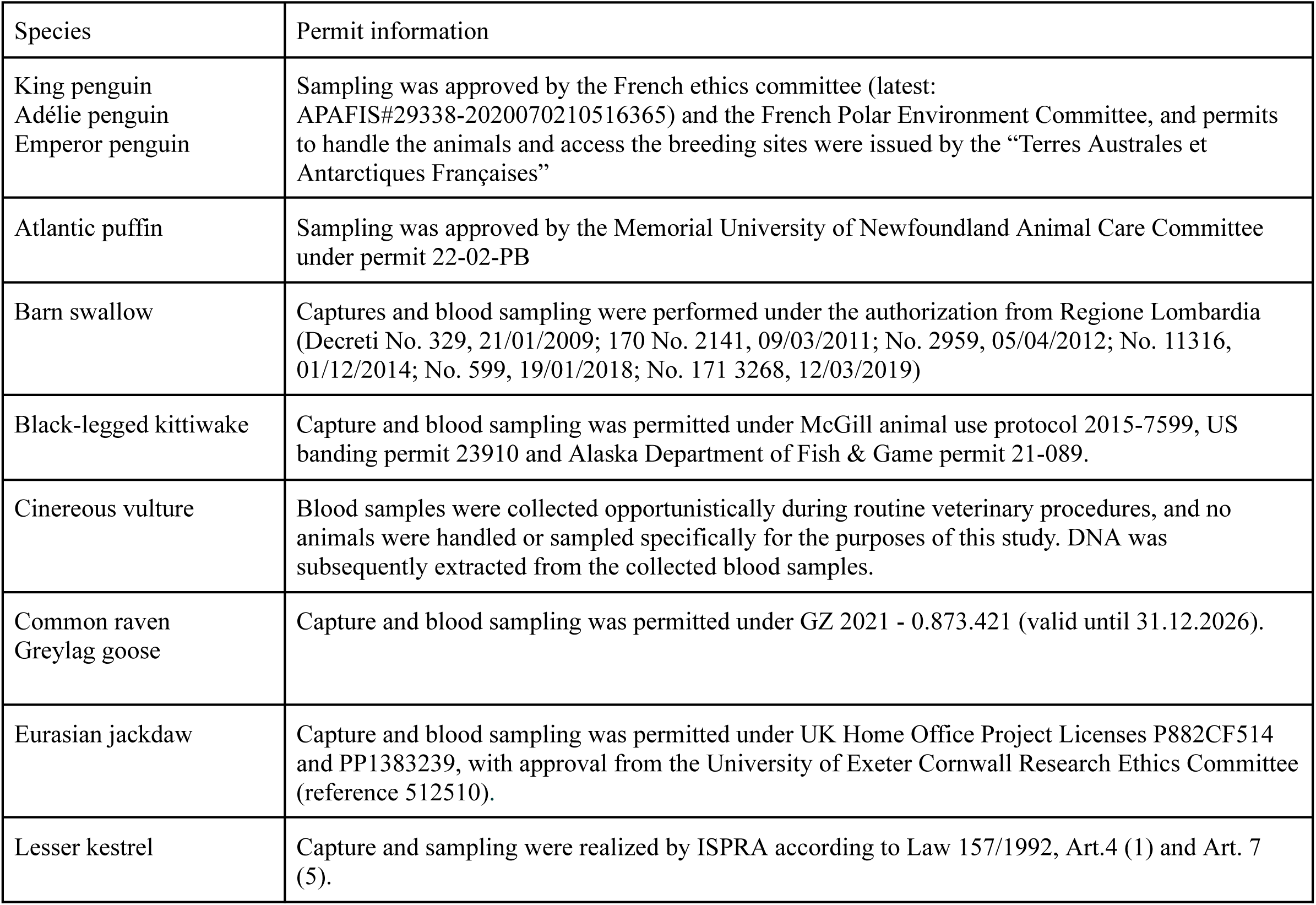
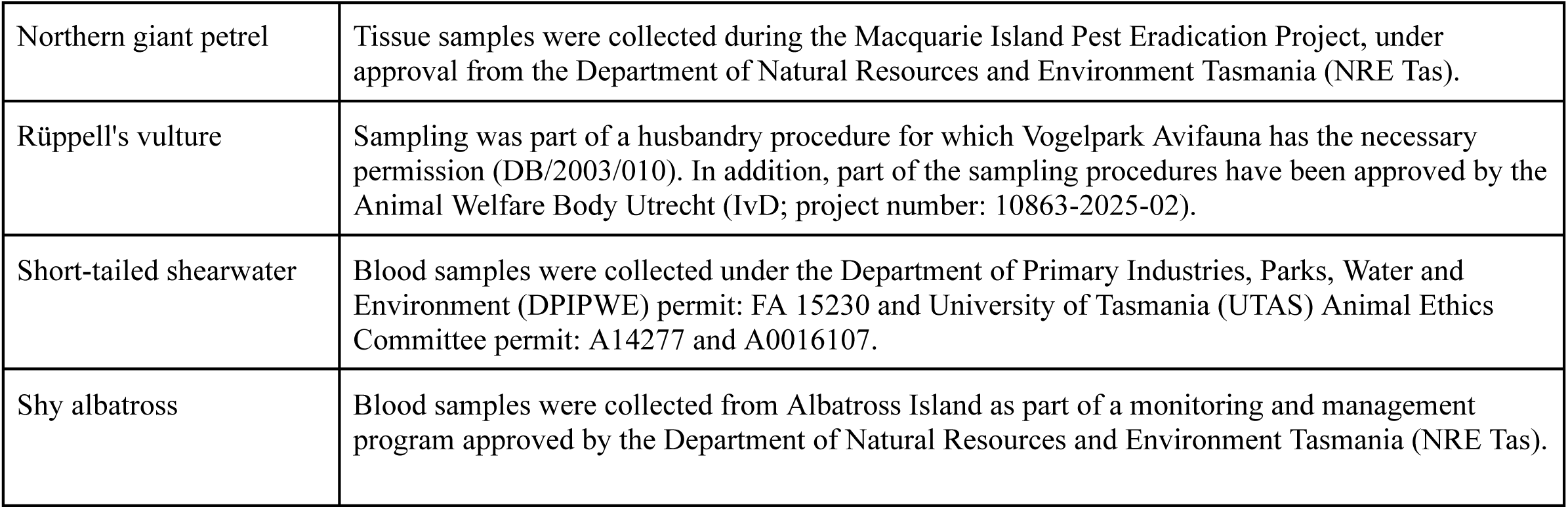

| Species | Permit information |
| --- | --- |
| King penguin<br>Adélie penguin<br>Emperor penguin | Sampling was approved by the French ethics committee (latest: APAFIS#29338-2020070210516365) and the French Polar Environment Committee, and permits to handle the animals and access the breeding sites were issued by the “Terres Australes et Antarctiques Françaises” |
| Atlantic puffin | Sampling was approved by the Memorial University of Newfoundland Animal Care Committee under permit 22-02-PB |
| Barn swallow | Captures and blood sampling were performed under the authorization from Regione Lombardia (Decreto No. 329, 21/01/2009; 170 No. 2141, 09/03/2011; No. 2959, 05/04/2012; No. 11316, 01/12/2014; No. 599, 19/01/2018; No. 171 3268, 12/03/2019) |
| Black-legged kittiwake | Capture and blood sampling was permitted under McGill animal use protocol 2015-7599, US banding permit 23910 and Alaska Department of Fish & Game permit 21-089. |
| Cinereous vulture | Blood samples were collected opportunistically during routine veterinary procedures, and no animals were handled or sampled specifically for the purposes of this study. DNA was subsequently extracted from the collected blood samples. |
| Common raven<br>Greylag goose | Capture and blood sampling was permitted under GZ 2021 - 0.873.421 (valid until 31.12.2026). |
| Eurasian jackdaw | Capture and blood sampling was permitted under UK Home Office Project Licenses P882CF514 and PP1383239, with approval from the University of Exeter Cornwall Research Ethics Committee (reference 512510). |
| Lesser kestrel | Capture and sampling were realized by ISPRA according to Law 157/1992, Art.4 (1) and Art. 7 (5). |
| Northern giant petrel | Tissue samples were collected during the Macquarie Island Pest Eradication Project, under approval from the Department of Natural Resources and Environment Tasmania (NRE Tas). |
| Rüppell's vulture | Sampling was part of a husbandry procedure for which Vogelpark Avifauna has the necessary permission (DB/2003/010). In addition, part of the sampling procedures have been approved by the Animal Welfare Body Utrecht (IvD; project number: 10863-2025-02). |
| Short-tailed shearwater | Blood samples were collected under the Department of Primary Industries, Parks, Water and Environment (DPIPWE) permit: FA 15230 and University of Tasmania (UTAS) Animal Ethics Committee permit: A14277 and A0016107. |
| Shy albatross | Blood samples were collected from Albatross Island as part of a monitoring and management program approved by the Department of Natural Resources and Environment Tasmania (NRE Tas). |

## Institutional and computational resources

We acknowledge the Competence Centre for Genomic Analysis (Kiel) for sequencing the samples of the EM-Seq data, and the DNA Sequencing and Genomics Laboratory (supported by HiLIFE and Biocenter Finland funding), Institute of Biotechnology, University of Helsinki for sequencing of the BEAC-Seq data. The authors wish to further acknowledge CSC – IT Center for Science, Finland, for computational resources, the DFG Research Infrastructure NGS_CC (project 407495230) as part of the Next Generation Sequencing Competence Network (project 423957469), the Department of Work, Economy, Science, Innovation and Social Economy (WEWIS) of the Flemish government for supporting the cinereous vulture sampling project, and the CSIRO Environomics Future Science Platform, Australia, for supporting OB, AB, and CA during the project. Figure 1 was created in BioRender.com

## People

We are deeply grateful to all the wintering and summering members of IPEV Project 137 and all the other colleagues and students within the P137 team, who participated in the king penguin long-term monitoring and sample collection since 2000. We sincerely thank the IPEV logistics teams for their important and continued support in the field. This study is part of and supported by the long-term Studies in Ecology and Evolution (SEE-Life) program of the CNRS. For collecting samples from Atlantic puffins we thank Maarten Vis of the Rotterdam Zoo, Gheylen Daghfous of the Biodôme de Montréal, Ana Ferreira of the Oceanário de Lisboa, Stephan Hoby of the Bern Animal Park, and Susanne Leitinger of the Loro Park. We further wish to acknowledge Josef Hemetsberger for his invaluable support in collecting greylag goose samples. We also thank Scott Hatch for his help in collecting known-age kittiwake samples. Finally, we thank the team members who collected samples across various bird species not explicitly mentioned here; their work expanded this study’s scope, making it a valuable resource for ornithologists and researchers beyond the king penguin community.

## Author contributions

The study was designed by MH, RC, MO, SJ, BSM, and CLB. Sample contribution was done by ACC, AC, AR, AB, AT, AM, AP, AP, DJLB, DF, EVD, EP, FANF, GEM, GB, JL, JP, JFO, JGC, JM, KHE, LRD, LR, MH, MV, MM, MC, NC, NCS, NG, OB, PPB, PP, PS, PH, RC, RA, SJ, SK, SH, SW, TB, CT, and ET. Laboratory work was performed by MH, CA, ME, SW, AP, AS, SZ, NCS and LP. Data preparation and analysis was designed and conducted by MH, FNF, SJ, MO and RC. The first draft of the manuscript was written by MH, and all authors commented on the manuscript. All authors read and approved the final manuscript. MO and RC jointly supervised this work.

