## Supplementary Information for "The BEAC, an epigenetic clock for birds"

### Supplementary Tables

**Supplementary Table 1. Bird samples used in the analyses of the manuscript.** Samples used in different analysis steps with their corresponding distribution of species, DNA methylation measurement method (Method), number of individuals (Ind) and sample sizes (Sample), age range in years (Age), precision of age estimation (Precision), sex (F=female, M=male, NA), location of sampling, and rearing environment (Z=zoo-housed, W=wild, NA).

| Step | Species | Method | Ind | Sample | Age | Precision | F/M/NA | Location | Z/W/NA |
|---|---|---|---|---|---|---|---|---|---|
| Discovery | King penguin | EM-Seq | 87 | 96 | 0.88–35.20 | Month | 30/57 | Crozet/Zoo Zürich/Loro Parque | 30/57/0 |
| Training | King penguin | EM-Seq | 78 | 78 | 0.88–23.12 | Month | 36/42/0 | Crozet | 0/78/0 |
| Testing | King penguin | BEAC | 41 | 41 | 0.07-22.0 | Month | 20/15/6 | Crozet | 0/41/0 |
| Longitudinal | King penguin | BEAC | 7 | 17 | 0.9-19 | Month | 2/5/0 | Crozet | 0/7/0 |
| Multispecies | | | | | | | | | |
| | Adélie penguin | BEAC | 11 | 12 | 0-20 | Year | 0/0/12 | Adélie land, Antarctica | 0/12/0 |
| | Atlantic puffin | BEAC | 15 | 15 | 0.7- 32.8 | Day | 7/6/2 | Biodôme/Bern Animal Park/Rotterdam Zoo/Loro Park/Oceanário de Lisboa | 15/0/0 |
| | Barn swallow | BEAC | 20 | 20 | 0-8 | Year | 7/13/0 | Milan, Italy | 0/20/0 |
| | Black-legged kittiwake | BEAC | 20 | 20 | 0-21 | Year | 8/12/0 | Middleton Island, Alaska, USA | 0/20/0 |
| | Cinereous vulture | BEAC | 18 | 18 | 0-36 | Day | 11/7/0 | EAZA (European Association of Zoos and Aquaria)*[1] | 18/0/0 |
| | Common raven | BEAC | 2 | 2 | 5.46-8.51 | Day | 0/2/0 | Alm Valley, Austria | 2/2/0*[2] |
| | Emperor penguin | BEAC | 2 | 2 | 0 | Month | 0/0/2 | Adélie land, Antarctica | 0/2/0 |
| | Greylag goose | BEAC | 10 | 10 | 1-18 | Year | 1/8/1 | Alm Valley, Austria | 0/10/0 |
| | Eurasian jackdaw | BEAC | 17 | 17 | 0-12 | Year | 8/6/3 | Cornwall, England | 0/17/0 |
| | Lesser kestrel | BEAC | 18 | 18 | 0-9 | Year | 8/10/0 | Matera, Italy | 0/18/0 |
| | Northern giant petrel*[3] | BEAC | 11 | 11 | 4-22 | Year | 5/6/0 | Macquarie Island, Tasmania, AU | 0/11/0 |
| | Rüppell's vulture | BEAC | 11 | 11 | 1.03-16.05 | Day | 7/4/0 | Avifauna Bird Park, Netherlands | 11/0/0 |
| | Short-tailed shearwater | BEAC | 12 | 12 | 0.15-19 | Year | 4/7/1 | Flinders Island, Tasmania, AU | 0/12/0 |
| | Shy albatross | BEAC | 11 | 11 | 4-18 | Year | 6/5/0 | Albatross Island, Tasmania, AU | 0/11/0 |

*1 Planckendael, Belgium; Pairi Daiza, Cambron, Belgium; Haringsee, Austria; Landskron, Austria; Les Eppesses Puy du Fou, France; Mulhouse, France; Puy d Fou España, Spain; Tallinn, Estonia; Green Balkans, Bulgaria; Liberec, Czech Republic

*2 Raised under human care for ~6 months, and released into the wild

*3 Muscle samples (all other samples are blood)

**Supplementary Table 2. BEAC primer sequences for King penguins.** Primer pair, sequences for forward and reverse primers, PCR pool and amplicon size are given for each of the 24 primer pairs.

| Pair | Forward | Reverse | Pool | Size |
|---|---|---|---|---|
| H9 | GTGGGTAGAGGGTATGGT | AACTTAACAACCTACTACATCC | A | 131 |
| H25 | TATCAATAACTTATCACTTACCTC | AGATGTTTGGTATGGTTTTGTG | A | 118 |
| H34 | AGTTAATTTGGTAGTTAGTAATTGAT | AATCTTAACCCAAATCTACCTC | A | 100 |
| H42 | TCCAAACATTTAATACCTAATTAAATA | TGGAATATATAGGGTTTGGAAG | A | 112 |
| H60 | CTACACAAAAACTTTAAACACACA | AGGTTGATGTTTAGAGATTATAGA | A | 131 |
| H62 | TACATATTAATATACCACACAACC | AGTTTGTTTGATAAAGATGGATATAA | A | 130 |
| H73 | CCATTAAACAAAAATTAACACTATTC | GTAAATGATTGAAAGTGTTTTTAGAT | A | 134 |
| H79 | TTTCCCAACACCTCTTAACAAT | TTGGGGGTAGAAATGTGAGA | A | 143 |
| H103 | ATACTACCTATACTCTAAAAAACC | GTTGTGATTTAAAAGTTGGGGA | A | 137 |
| H116 | TGTAGAGTAGTGGTTTGTAGG | AACAAAAAACCACACCTAACTTC | A | 122 |
| H126 | AAAAAAACAATCCAACAAAAATCACA | TATGTATATAAAGAAGAAAAGTTGAG | A | 120 |
| H128 | AAGGTTTGGTTATGGTTTGTGT | CAAACTATTTAATACCATAATCCC | A | 108 |
| H12 | ATATTAAAATATACACCCCTACCA | GTTTAGTATAGGTTTTGTTTGAGT | B | 120 |
| H16 | CTCCAACCCATCCTACCT | TAAGGGGATGTAGTTTTTGTGT | B | 115 |
| H26 | CTTAAAAACACTTAACTTACAATATAA | TTTAGGGGGAGTATTTTGGG | B | 127 |
| H32 | TAAATTTCTCTATTTACCATAATAATTC | TTTGGTGAGGTAGAGGGATA | B | 125 |
| H102 | TACCCTAAAACTTCTTACCAAC | TTAGGTGAGTTTTTTTTAATAATAGG | B | 145 |
| H104 | CCTACAAACCCTAAAAAAAATCC | TTTGGTTAGAAGTTTAGATGTTGA | B | 110 |
| H106 | GGTTATTTAATTGTTATTTTTGTAGTT | TCTTATTACCTAAAACCCTTACTT | B | 147 |
| H119 | GTTGTTTTAGTATGTGGTGGAA | ACCAAACAAACACAAACAATATCA | B | 100 |
| H125b | GGTTTAAGAAGAATTGTTGGGT | CTCAAATTCTTTAATCTAAATATTTCC | B | 114 |
| H129 | GTGGTATTGTTGGAGTGATG | AACCATTTACAACATATCCCAC | B | 129 |
| H131 | CACCTAACAAAACACCCCAT | TTATTGGTAGAGTTTTTTGTTGGA | B | 123 |
| H135 | AACTCCCATCATCTCAATCC | GGTATTTGAGTGTTGTGGTTAT | B | 145 |

**Supplementary Table 3. BEAC primer sequences for additional bird species.** Names and sequences of primers recommended for use in avian species outside of King penguins.

| Name | Sequence |
|---|---|
| H16Fa1 | CTCCAACCCATCCCRCCT |
| H32Fa1 | TAAATTTCTCTATTTACCACRATAATTC |
| H32Ra1 | TTTGGTGGGGTAGAGGGATA |
| H34Fa1 | AGYGAATTTGGTAGTTAGTAATTGAT |
| H34Fa2 | AGTTAATTTGGTAGTTAGTAATTGAG |
| H62Fa1 | TACRTATTACTATACCACACAACC |
| H62Fa2 | TACATATTACTATACCACAAAACC |
| H62Ra1 | AGTTTGTTTGATAAAGATAGATATAA |
| H62Ra2 | AGTTTGTTTGATTAAGATAGATATAA |
| H73Fa1 | CCRTTAAACAAAAATTAACACTATTC |
| H73Ra1 | GYGAATGATTGAAAGTGTTTTTAGAT |
| H79Fa1 | TTTTCCCAACACCTCTTAAC |
| H79Ra1 | TTGGGGGTAGAAATGTGAGC |
| H79Ra2 | TYGGGGGTAGAAATGTGAG |
| H79Ra3 | TTTGGGGGTAGAAATGTGAG |
| H104Fa1 | CCTACAATCCTTATAAAAAATCC |
| H104Fa2 | CCTACAAAACCTAAAAAAAATCC |
| H104Fa3 | CCTACAATCTTTATAAAAAATCC |
| H104Ra1 | TTTGGTTAGGAGTTTAGATGTTGA |
| H104RY | TTTGGTTAGAAGTTTAGAYGTTGA |
| H125bFa3 | GGTTTAAGAAAAATTGTTGGGT |
| H125bFa4 | GGTTTAAGAAAAATTGTTAGGT |
| H131Fa1 | CATCTAACAAAACATCCCAT |
| H131Fa2 | CRTCTATCAAAACACCCCRT |
| H131Fa3 | CATCTAACAAAACATCACAT |
| H131Ra2 | TTATTGGTAGAGTTTTTTGTTGTT |
| H131Ra3 | TTATTGTTAGAATTTTTTGTTGGA |

**Supplementary Table 4. Reference species used in primer design and sequencing data alignment for the sampled species.** Primer design reference represents the species whose genome was used in primer design retrieved from multispecies alignment published in Feng et al. 2020, incorporated with the King penguin genome. Alignment reference corresponds to the species used in sequencing data alignment with the corresponding accession code.

| Sampled species | Primer design reference | Alignment reference | Accession |
|---|---|---|---|
| King penguin (*Aptenodytes patagonicus*) | King penguin (*Aptenodytes patagonicus*) | King penguin (*Aptenodytes patagonicus*) | GCA_010087175.1 |
| Adélie penguin (*Pygoscelis adeliae*) | Adélie penguin (*Pygoscelis adeliae*) | Adélie penguin (*Pygoscelis adeliae*) | GCF_000699105.1 |
| Emperor penguin (*Aptenodytes forsteri*) | Emperor penguin (*Aptenodytes forsteri*) | Emperor penguin (*Aptenodytes forsteri*) | GCF_000699145.1 |
| Eurasian jackdaw (*Coloeus monedula*) | Hooded crow (*Corvus cornix*) | Eurasian jackdaw (*Coloeus monedula*) | GCF_965178545.1 |
| Barn Swallow (*Hirundo rustica*) | Barn Swallow (*Hirundo rustica*) | Barn Swallow (*Hirundo rustica*) | GCF_015227805.2 |
| Black-legged kittiwake (*Rissa tridactyla*) | Black-legged kittiwake (*Rissa tridactyla*) | Black-legged kittiwake (*Rissa tridactyla*) | GCF_028500815.1 |
| Lesser kestrel (*Falco naumanni*) | Peregrine falcon (*Falco peregrinus*) | Lesser kestrel (*Falco naumanni*) | GCF_017639655.2 |
| Rüppell's vulture (*Gyps rueppelli*) | Black-chested snake eagle (*Circaetus pectoralis*) | Rüppell's vulture (*Gyps rueppelli*) | GCA_027575145.1 |
| Cinereous vulture (*Aegypius monachus*) | Black-chested snake eagle (*Circaetus pectoralis*) | Cinereous vulture (*Aegypius monachus*) | GCA_982185325.1 |
| Greylag goose (*Anser anser*) | Swan goose (*Anser cygnoides*) | Greylag goose (*Anser anser*) | GCF_964211835.1 |
| Atlantic puffin (*Fratercula arctica*) | Razorbill (*Alca torda*) | Atlantic puffin (*Fratercula arctica*) | GCA_947846985.1 |
| Northern giant petrel (*Macronectes halli*) | Northern fulmar (*Fulmarus glacialis*) | Northern giant petrel (*Macronectes halli*) | GCA_977968185.1 |
| Shy albatross (*Thalassarche cauta*) | Atlantic yellow-nosed albatross (*Thalassarche chlororhynchos*) | Shy albatross (*Thalassarche cauta*) | GCA_013400895.1 |
| Short-tailed shearwater (*Ardenna tenuirostris*) | Cory's shearwater (*Calonectris borealis*) | Great shearwater (*Ardenna gravis*) | GCA_045784155.1 |

*Feng, S., Stiller, J., Deng, Y. et al. Dense sampling of bird diversity increases power of comparative genomics. Nature 587, 252–257 (2020).* https://doi.org/10.1038/s41586-020-2873

### Supplementary Figures

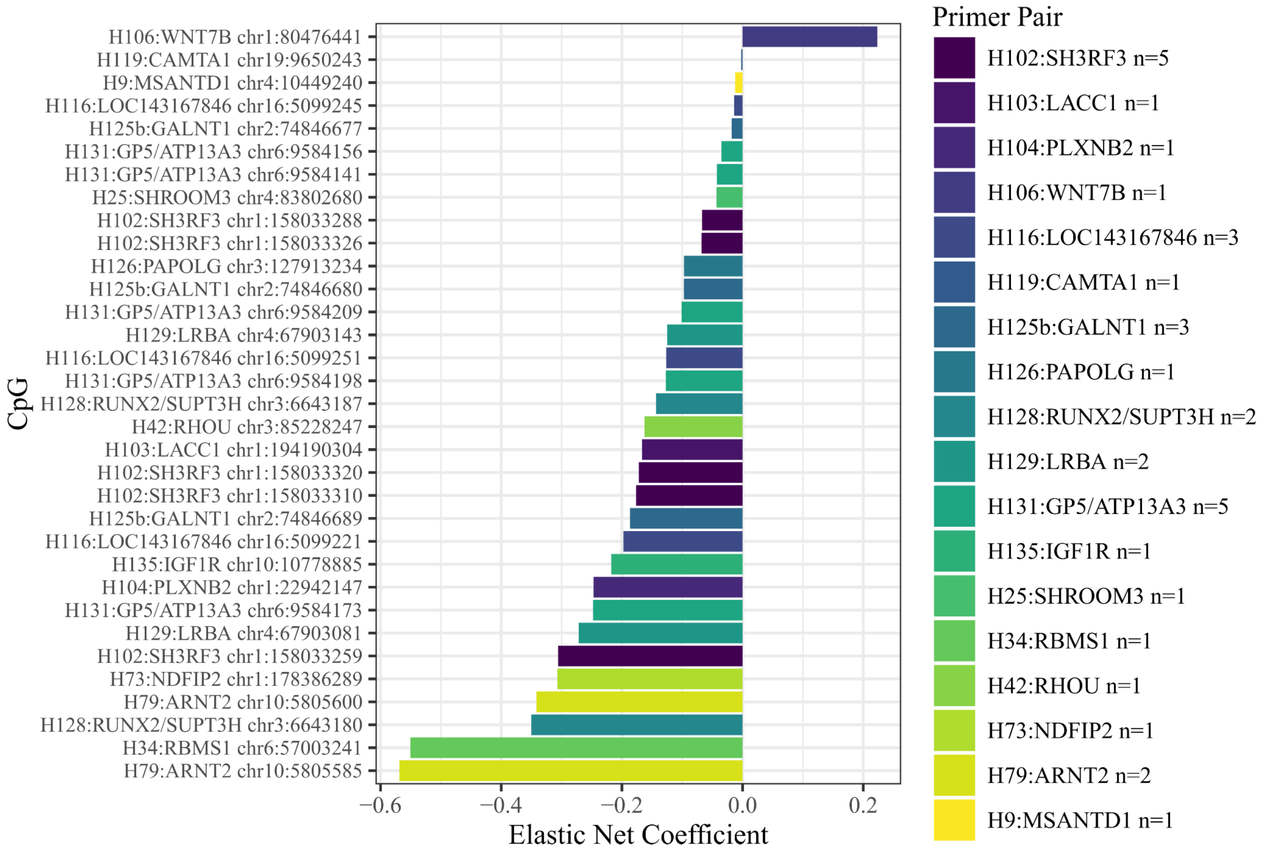


**Supplementary Figure 1. BEAC-Clock coefficients.** Model coefficients for each CpG selected by the elastic net model n(CpG)=33, n(Primer pair)=18. The number of predictors from each primer is listed on the right. Primer pairs are annotated by the closest gene of the amplicon.

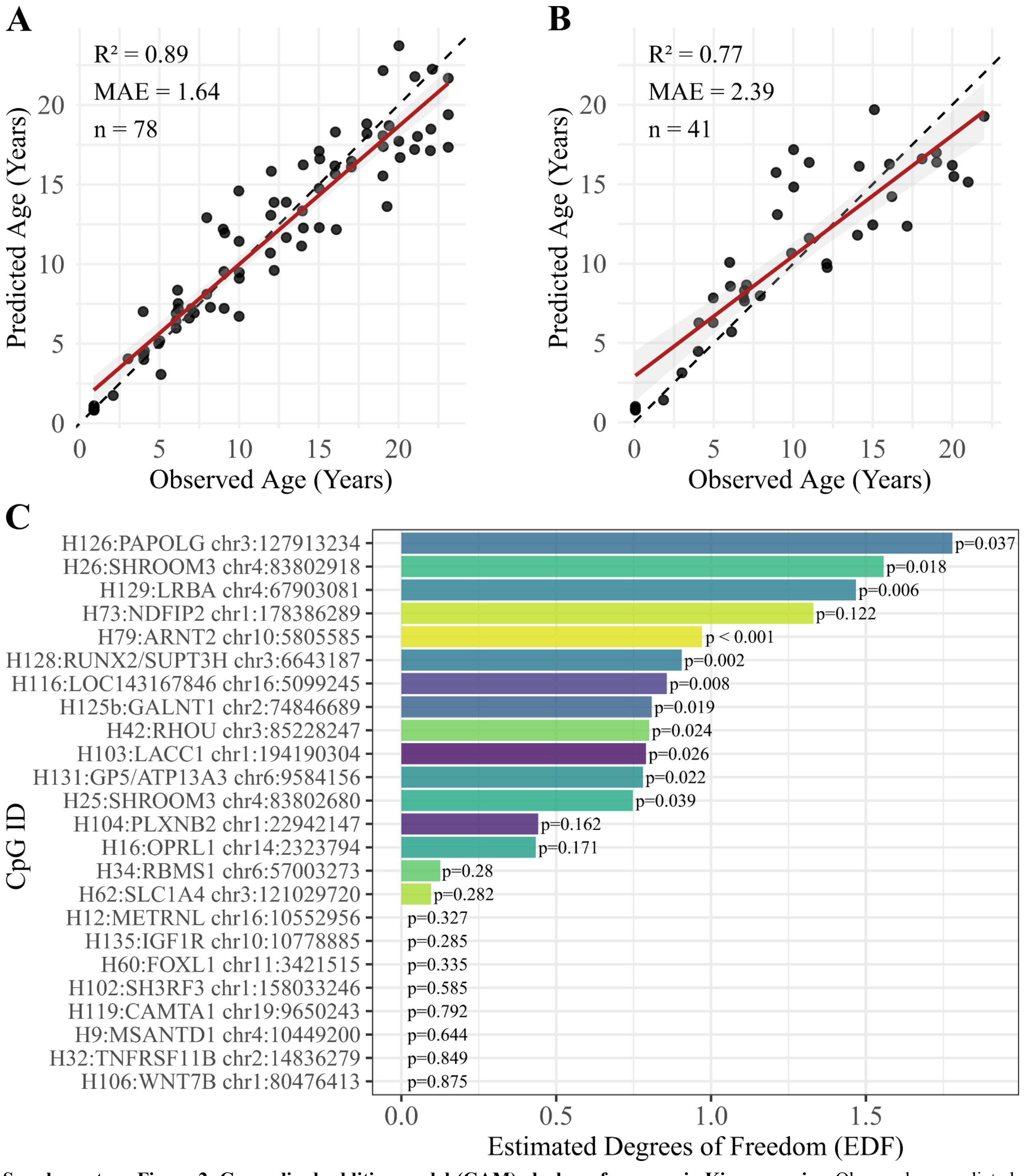


**Supplementary Figure 2. Generalised additive model (GAM)-clock performance in King penguins.** Observed vs. predicted age for the training data (A, n=78) and testing data (B, n=41). The dashed lines represent the 1:1 identity line. (C) Estimated effective degrees of freedom for each CpG predictor with associated p-values.

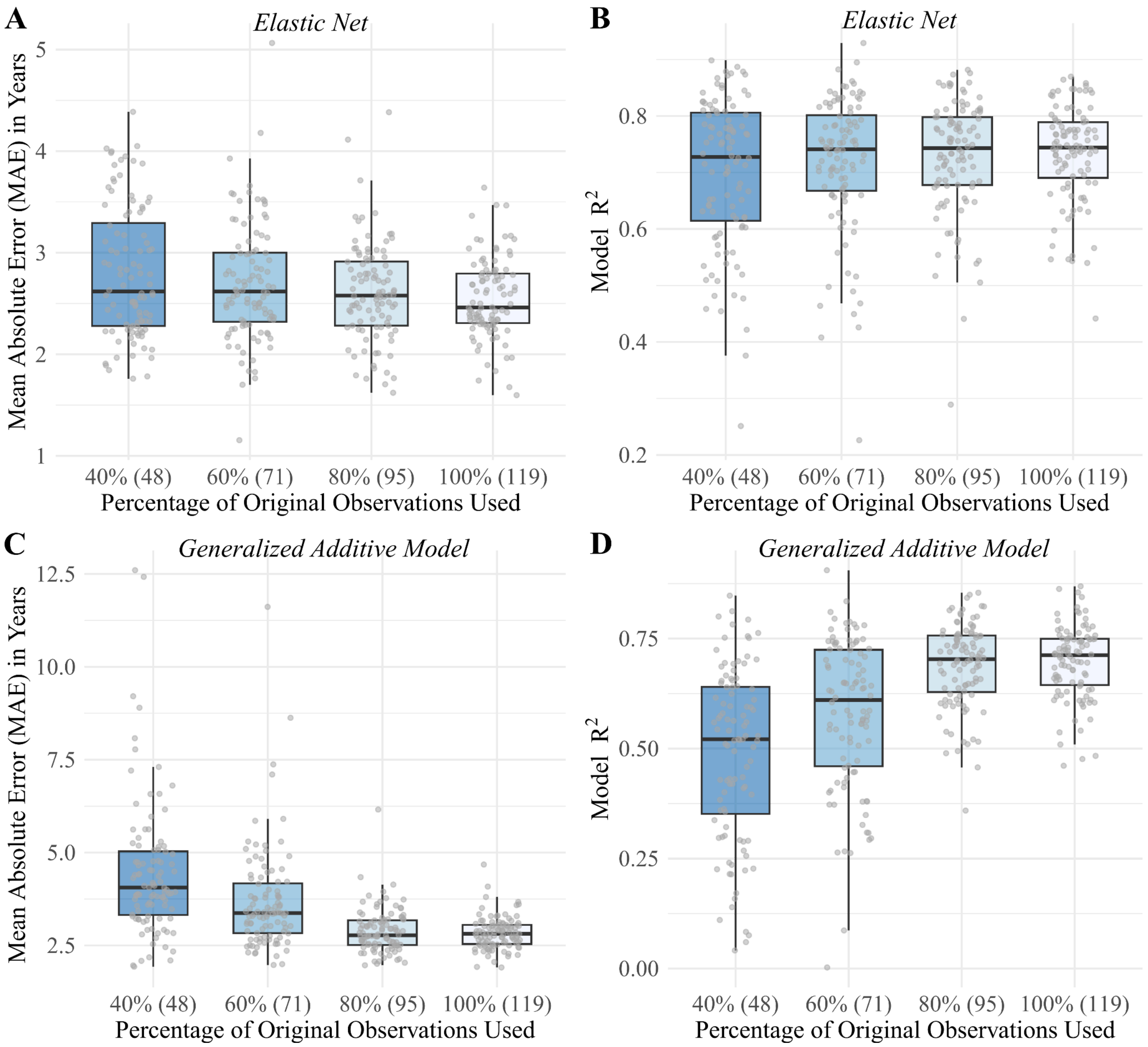


**Supplementary Figure 3. Model performance across varying training sample sizes.** To evaluate the impact of dataset scale on predictive accuracy, the initial dataset (n=119) was downsampled to 80%, 60%, and 40% of its original size. At each downsampling level, the data was randomly partitioned into training and testing sets over 100 iterations, and the models were retrained. Testing data **(A)** Mean absolute error (MAE, years) and **(B)** $R^2$ by downsampling level within the elastic net framework. Testing data **(C)** MAE (years) and **(D)** $R^2$ by downsampling level within the generalised additive model framework.

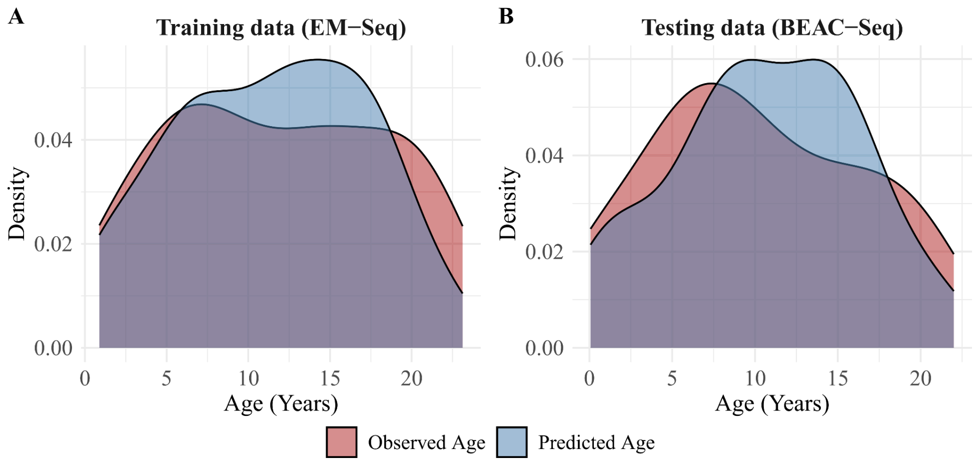


**Supplementary Figure 4. Distributions of observed and predicted age.** Overlaid density distributions of observed and predicted ages with the BEAC-Model in the training data (A, n=78) and testing data (B, n=41).

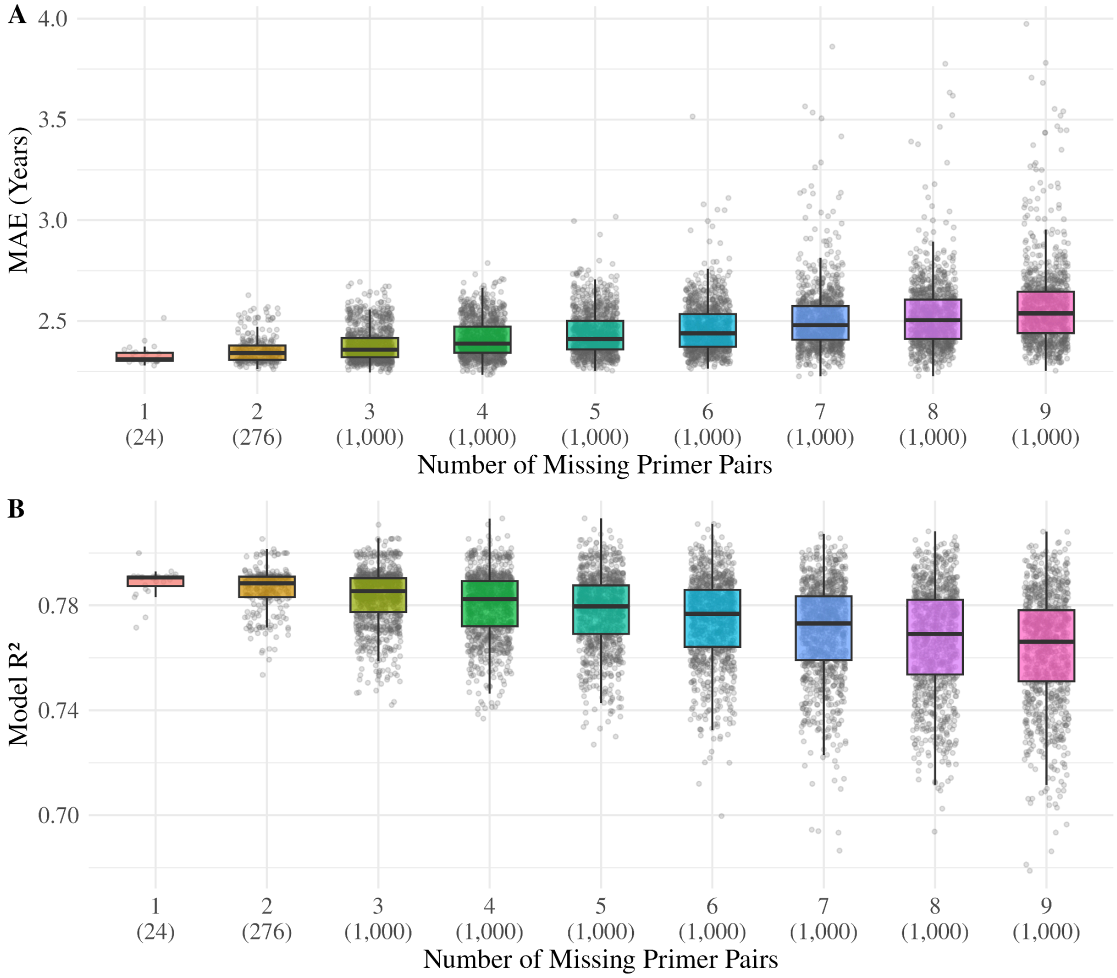


**Supplementary Figure 5. Impact of missing primer pairs on model performance.** Evaluation of 1–9 missing primer pairs of the BEAC-Seq across all possible missingness combinations, with missing values imputed using the training data panel; for counts yielding more than 100,000 possibilities, tested combinations were randomly subsampled and capped at 100,000. **(A)** Mean absolute error (MAE, years) and **(B)** $R^2$ relative to the number of excluded primer pairs. Values in parentheses indicate the total number of tested combinations for each count of missing pairs.

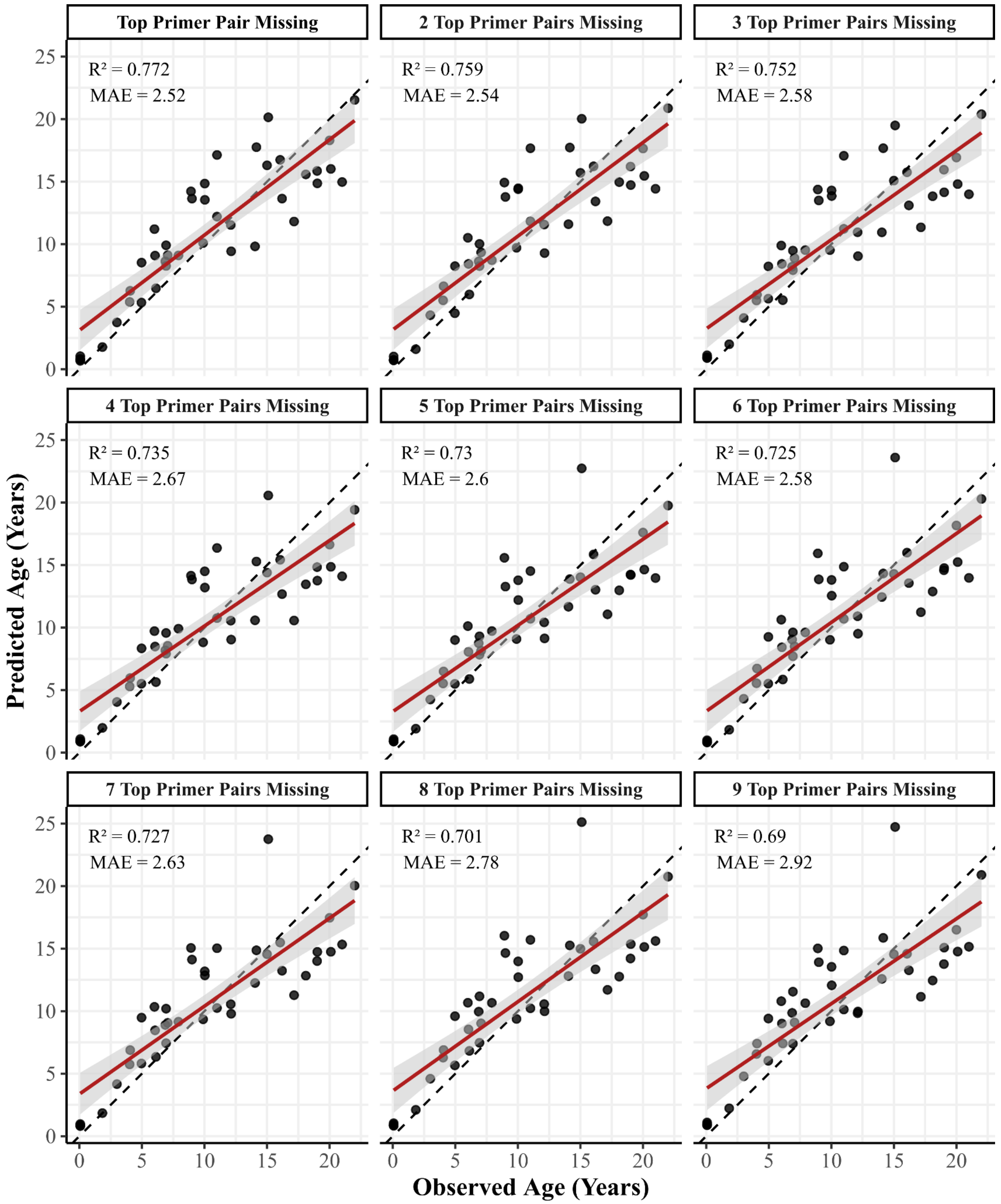


**Supplementary Figure 6. BEAC-Model performance with cumulative loss of the most influential amplicons.** Predicted vs. observed age in the independent testing dataset (n=41) following the iterative removal of top-ranked amplicons. Missing values were imputed via k-nearest neighbours using the training dataset. Amplicons were sequentially removed in descending order of importance (from the most important amplicon to the top nine ones), where importance was determined by ranking the total absolute sum of CpG coefficients within the BEAC-Model.

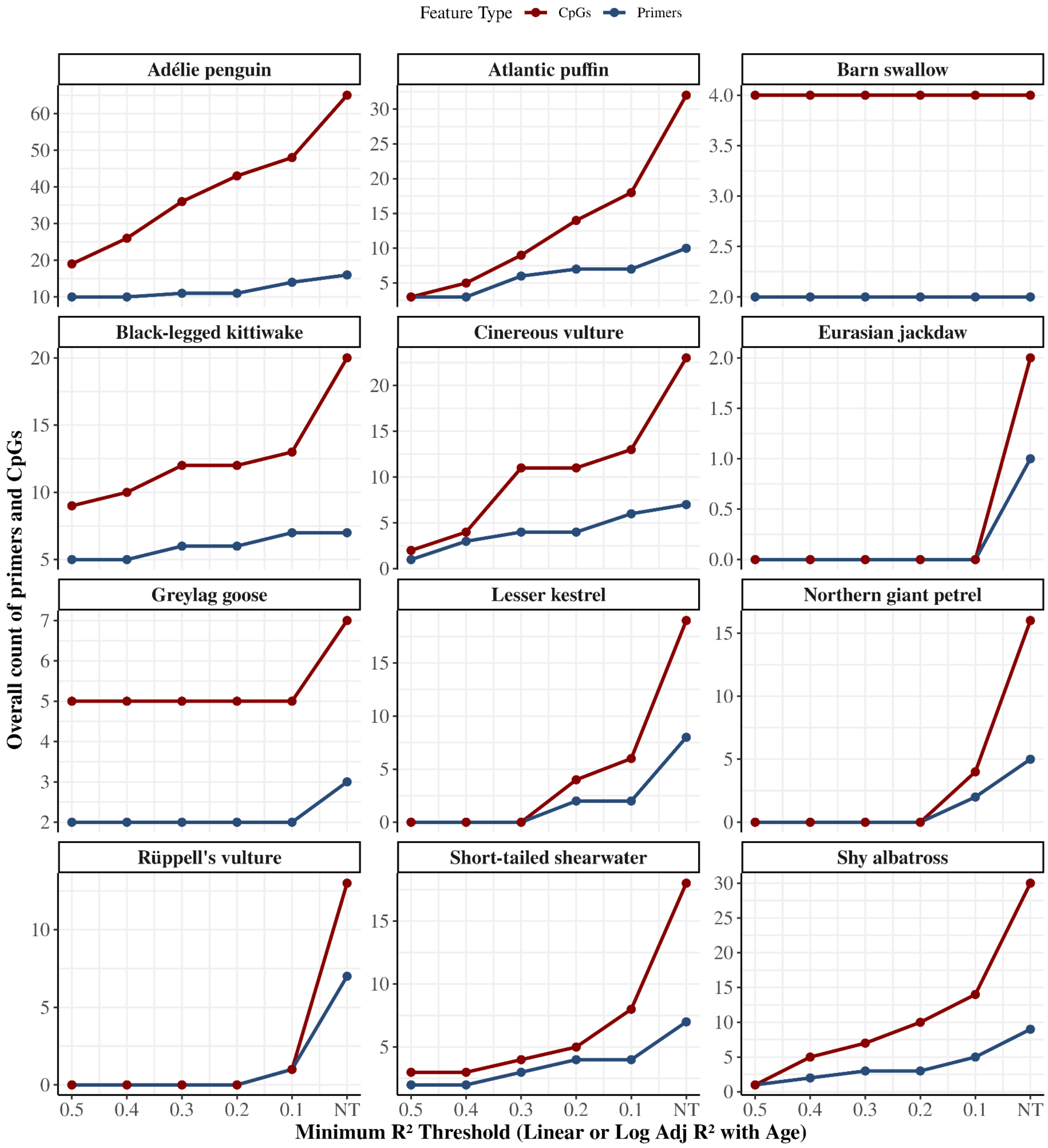


**Supplementary Figure 7. Feature counts across progressive $R^2$ filter thresholds by species.** The number of unique primers (blue) and individual CpG sites (red) across a sequence of minimum adjusted $R^2$. The adjusted $R^2$ is for the association between methylation and age at each CpG site (linear or log). The thresholds range from 0.5 to 0.1 and no threshold (NT).

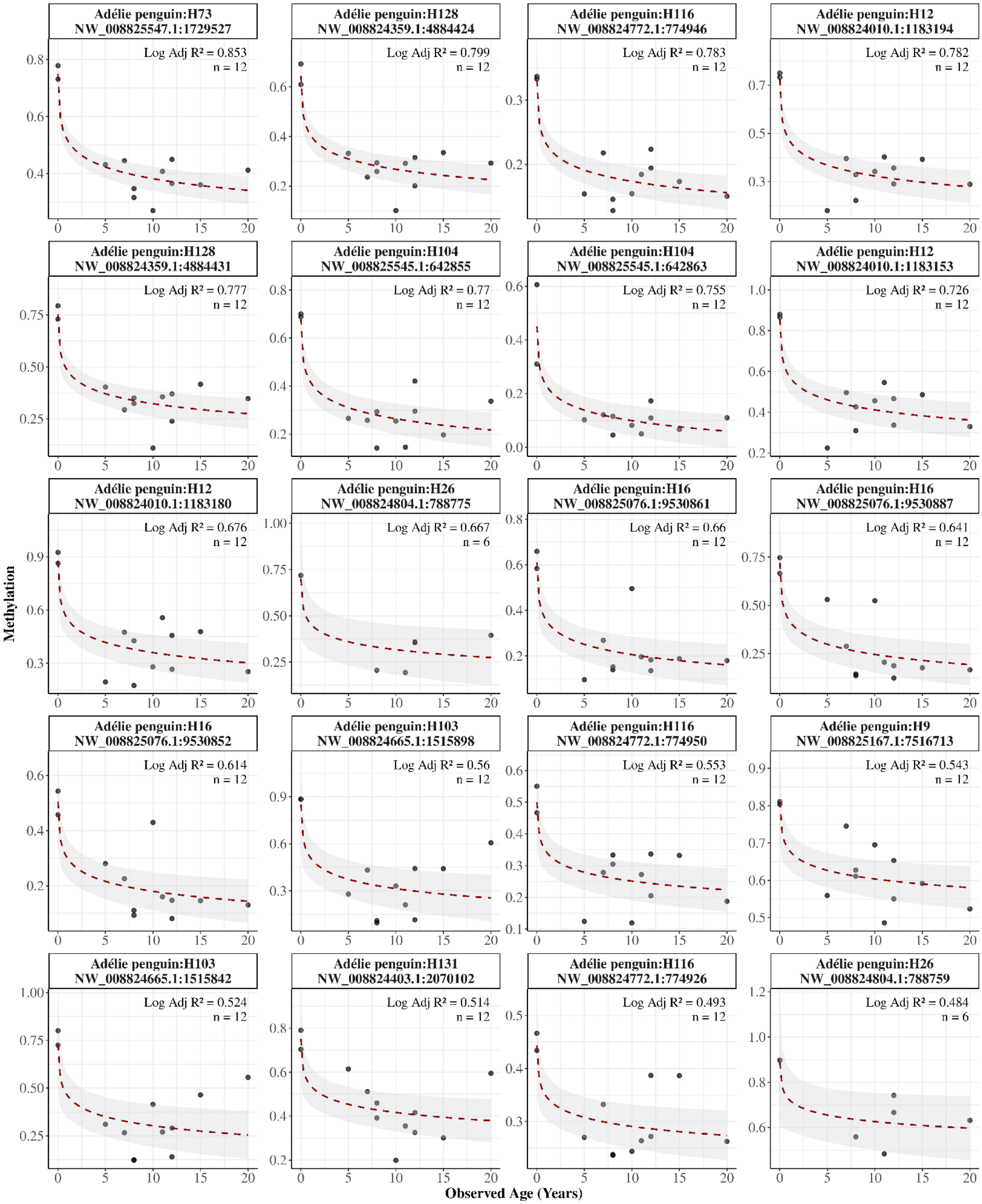

Adélie penguin:H73
NW_008825547.1:1729527
Log Adj R² = 0.853
n = 12
Adélie penguin:H128
NW_008824359.1:4884424
Log Adj R² = 0.799
n = 12
Adélie penguin:H116
NW_008824772.1:774946
Log Adj R² = 0.783
n = 12
Adélie penguin:H12
NW_008824010.1:1183194
Log Adj R² = 0.782
n = 12
Adélie penguin:H128
NW_008824359.1:4884431
Log Adj R² = 0.777
n = 12
Adélie penguin:H104
NW_008825545.1:642855
Log Adj R² = 0.77
n = 12
Adélie penguin:H104
NW_008825545.1:642863
Log Adj R² = 0.755
n = 12
Adélie penguin:H12
NW_008824010.1:1183153
Log Adj R² = 0.726
n = 12
Adélie penguin:H12
NW_008824010.1:1183180
Log Adj R² = 0.676
n = 12
Adélie penguin:H26
NW_008824804.1:788775
Log Adj R² = 0.667
n = 6
Adélie penguin:H16
NW_008825076.1:9530861
Log Adj R² = 0.66
n = 12
Adélie penguin:H16
NW_008825076.1:9530887
Log Adj R² = 0.641
n = 12
Adélie penguin:H16
NW_008825076.1:9530852
Log Adj R² = 0.614
n = 12
Adélie penguin:H103
NW_008824665.1:1515898
Log Adj R² = 0.56
n = 12
Adélie penguin:H116
NW_008824772.1:774950
Log Adj R² = 0.553
n = 12
Adélie penguin:H9
NW_008825167.1:7516713
Log Adj R² = 0.543
n = 12
Adélie penguin:H103
NW_008824665.1:1515842
Log Adj R² = 0.524
n = 12
Adélie penguin:H131
NW_008824403.1:2070102
Log Adj R² = 0.514
n = 12
Adélie penguin:H116
NW_008824772.1:774926
Log Adj R² = 0.493
n = 12
Adélie penguin:H26
NW_008824804.1:788759
Log Adj R² = 0.484
n = 6
Methylation
Observed Age (Years)

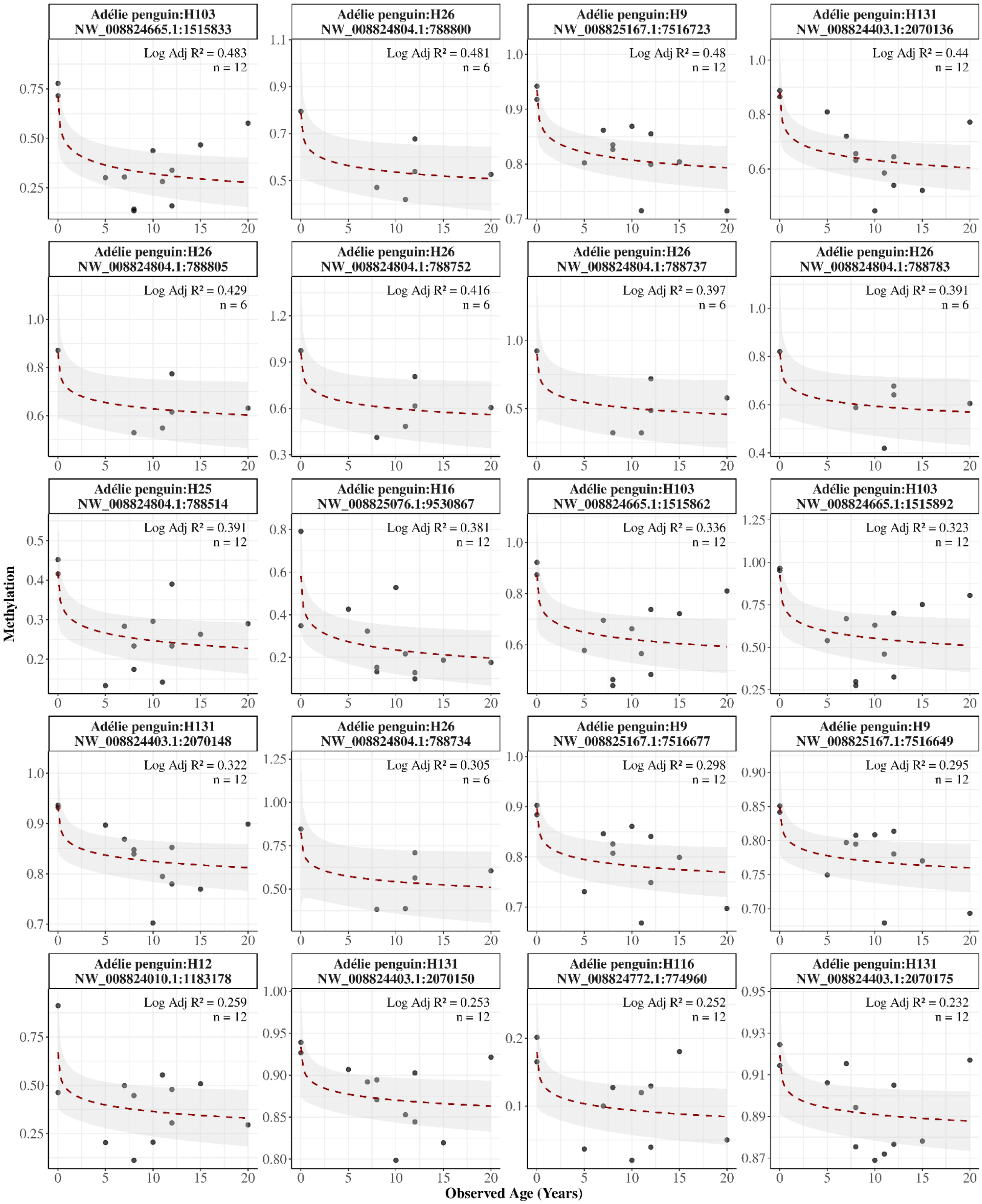
Adélie penguin:H103
NW_008824665.1:1515833
Log Adj R² = 0.483
n = 12
Adélie penguin:H26
NW_008824804.1:788800
Log Adj R² = 0.481
n = 6
Adélie penguin:H9
NW_008825167.1:7516723
Log Adj R² = 0.48
n = 12
Adélie penguin:H131
NW_008824403.1:2070136
Log Adj R² = 0.44
n = 12
Adélie penguin:H26
NW_008824804.1:788805
Log Adj R² = 0.429
n = 6
Adélie penguin:H26
NW_008824804.1:788752
Log Adj R² = 0.416
n = 6
Adélie penguin:H26
NW_008824804.1:788737
Log Adj R² = 0.397
n = 6
Adélie penguin:H26
NW_008824804.1:788783
Log Adj R² = 0.391
n = 6
Adélie penguin:H25
NW_008824804.1:788514
Log Adj R² = 0.391
n = 12
Adélie penguin:H16
NW_008825076.1:9530867
Log Adj R² = 0.381
n = 12
Adélie penguin:H103
NW_008824665.1:1515862
Log Adj R² = 0.336
n = 12
Adélie penguin:H103
NW_008824665.1:1515892
Log Adj R² = 0.323
n = 12
Adélie penguin:H131
NW_008824403.1:2070148
Log Adj R² = 0.322
n = 12
Adélie penguin:H26
NW_008824804.1:788734
Log Adj R² = 0.305
n = 6
Adélie penguin:H9
NW_008825167.1:7516677
Log Adj R² = 0.298
n = 12
Adélie penguin:H9
NW_008825167.1:7516649
Log Adj R² = 0.295
n = 12
Adélie penguin:H12
NW_008824010.1:1183178
Log Adj R² = 0.259
n = 12
Adélie penguin:H131
NW_008824403.1:2070150
Log Adj R² = 0.253
n = 12
Adélie penguin:H116
NW_008824772.1:774960
Log Adj R² = 0.252
n = 12
Adélie penguin:H131
NW_008824403.1:2070175
Log Adj R² = 0.232
n = 12
Methylation
Observed Age (Years)

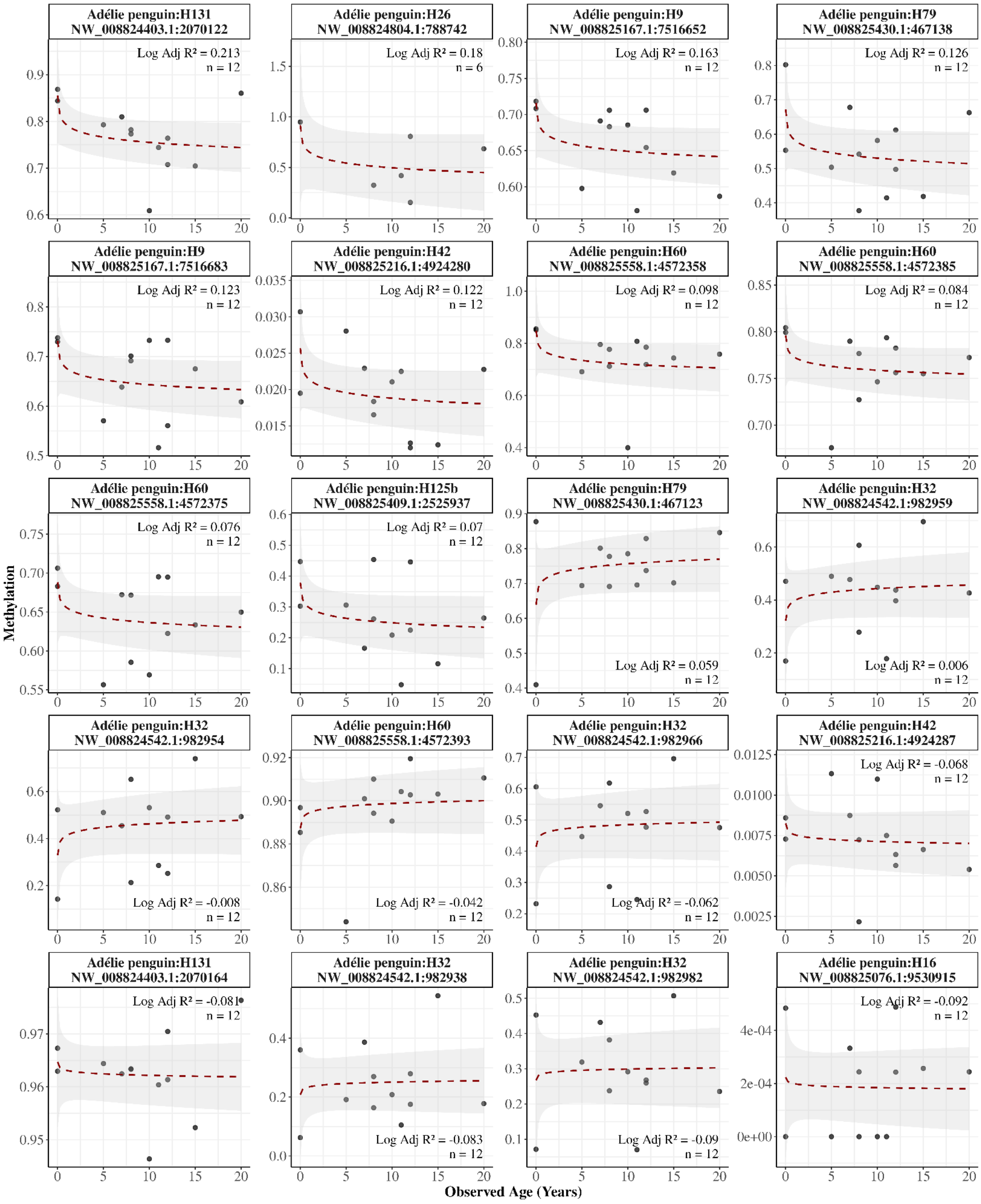
Adélie penguin:H131 NW_008824403.1:2070122
Log Adj R² = 0.213
n = 12
Adélie penguin:H26 NW_008824804.1:788742
Log Adj R² = 0.18
n = 6
Adélie penguin:H9 NW_008825167.1:7516652
Log Adj R² = 0.163
n = 12
Adélie penguin:H79 NW_008825430.1:467138
Log Adj R² = 0.126
n = 12
Adélie penguin:H9 NW_008825167.1:7516683
Log Adj R² = 0.123
n = 12
Adélie penguin:H42 NW_008825216.1:4924280
Log Adj R² = 0.122
n = 12
Adélie penguin:H60 NW_008825558.1:4572358
Log Adj R² = 0.098
n = 12
Adélie penguin:H60 NW_008825558.1:4572385
Log Adj R² = 0.084
n = 12
Adélie penguin:H60 NW_008825558.1:4572375
Log Adj R² = 0.076
n = 12
Adélie penguin:H125b NW_008825409.1:2525937
Log Adj R² = 0.07
n = 12
Adélie penguin:H79 NW_008825430.1:467123
Log Adj R² = 0.059
n = 12
Adélie penguin:H32 NW_008824542.1:982959
Log Adj R² = 0.006
n = 12
Adélie penguin:H32 NW_008824542.1:982954
Log Adj R² = -0.008
n = 12
Adélie penguin:H60 NW_008825558.1:4572393
Log Adj R² = -0.042
n = 12
Adélie penguin:H32 NW_008824542.1:982966
Log Adj R² = -0.062
n = 12
Adélie penguin:H42 NW_008825216.1:4924287
Log Adj R² = -0.068
n = 12
Adélie penguin:H131 NW_008824403.1:2070164
Log Adj R² = -0.081
n = 12
Adélie penguin:H32 NW_008824542.1:982938
Log Adj R² = -0.083
n = 12
Adélie penguin:H32 NW_008824542.1:982982
Log Adj R² = -0.09
n = 12
Adélie penguin:H16 NW_008825076.1:9530915
Log Adj R² = -0.092
n = 12
Methylation
Observed Age (Years)

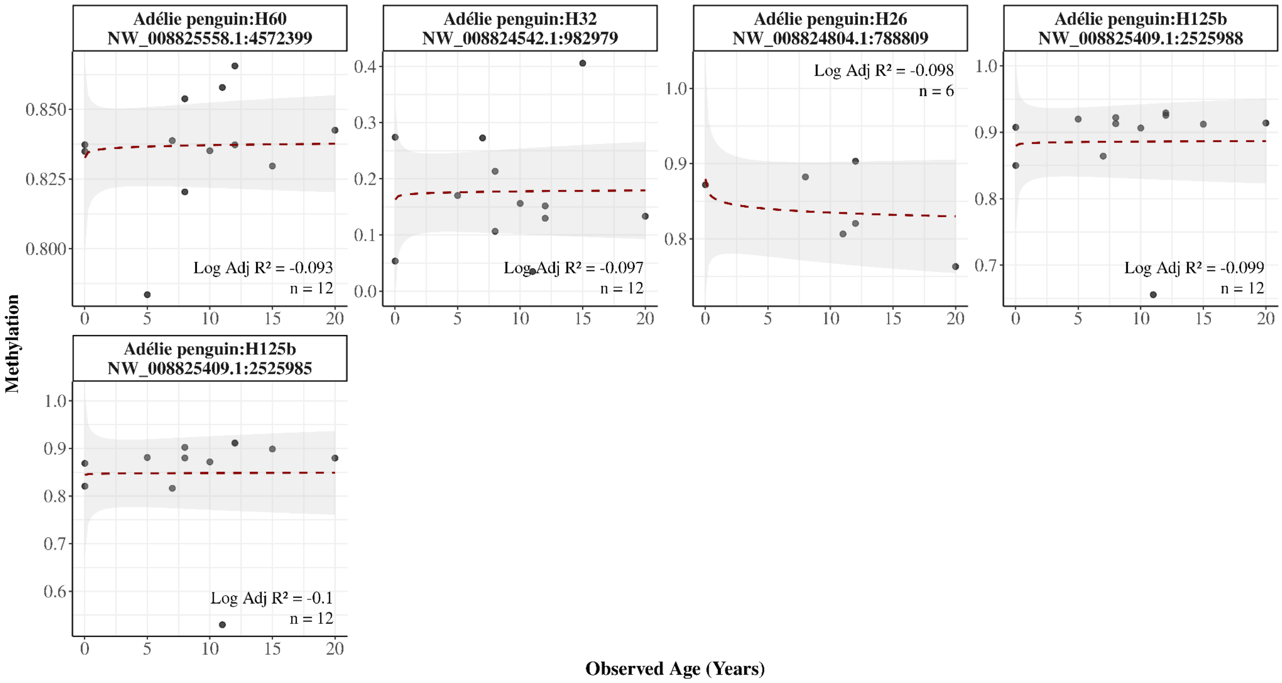


**Supplementary Figure 8. Age-dependent methylation in the Adélie penguin blood samples**. Panels display methylation levels (y-axis) against observed age (x-axis) for each CpG site ($n_{Sample}$=12, $n_{CpG}$=65, $n_{Primer}$=16). The plot includes all detected CpG sites with more than 3 samples, excluding those in which more than half of the samples showed extreme methylation values (exactly 0 or 1). Annotations include primer number, genomic position, logarithmic adjusted $R^2$ (Log Adj $R^2$) between methylation and age, and number of samples. CpG sites are ordered by logarithmic adjusted $R^2$, and a logarithmic smoothing line is applied to each panel.

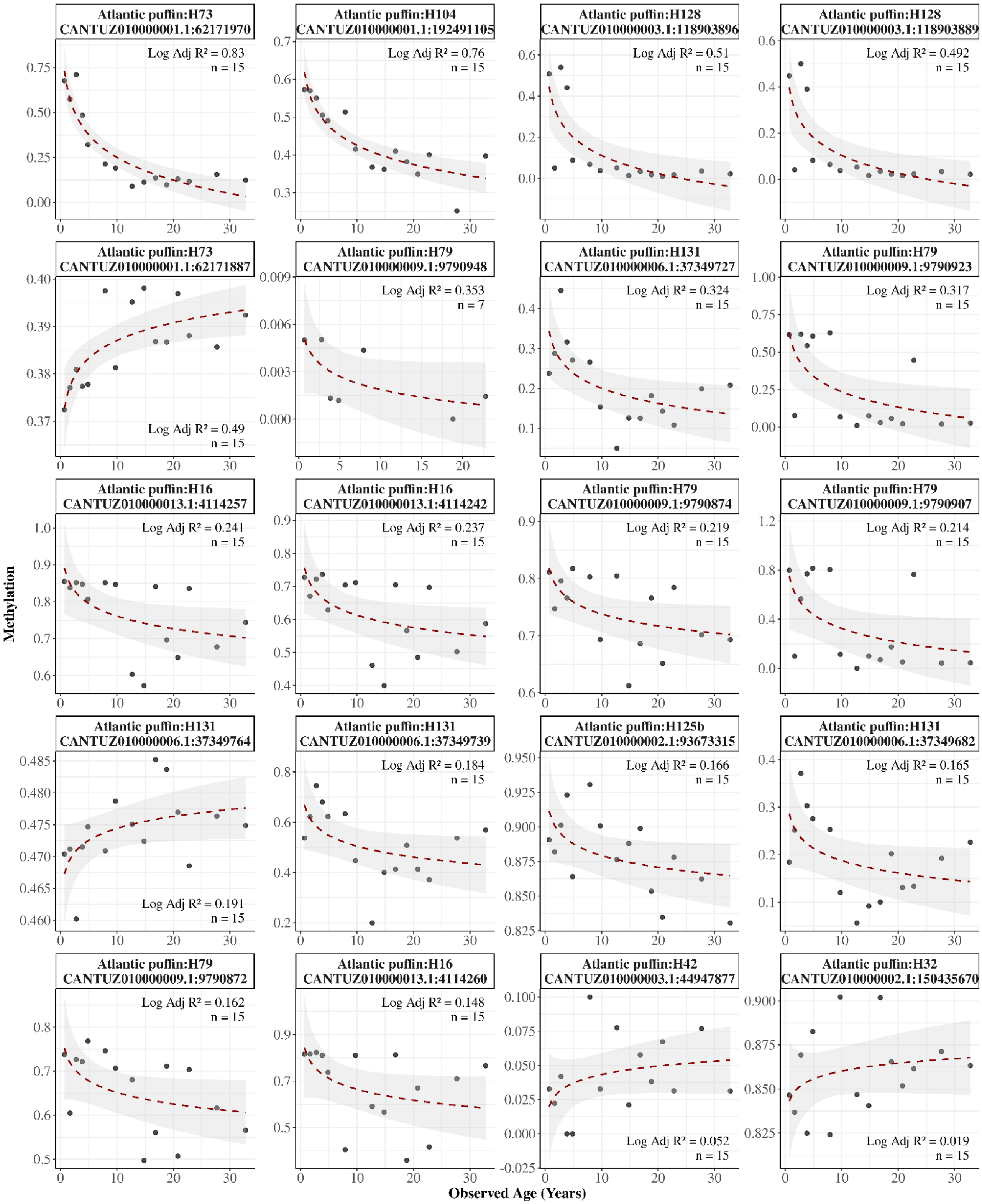

Atlantic puffin:H73
CANTUZ010000001.1:62171970
Log Adj R² = 0.83
n = 15
Atlantic puffin:H104
CANTUZ010000001.1:192491105
Log Adj R² = 0.76
n = 15
Atlantic puffin:H128
CANTUZ010000003.1:118903896
Log Adj R² = 0.51
n = 15
Atlantic puffin:H128
CANTUZ010000003.1:118903889
Log Adj R² = 0.492
n = 15
Atlantic puffin:H73
CANTUZ010000001.1:62171887
Log Adj R² = 0.49
n = 15
Atlantic puffin:H79
CANTUZ010000009.1:9790948
Log Adj R² = 0.353
n = 7
Atlantic puffin:H131
CANTUZ010000006.1:37349727
Log Adj R² = 0.324
n = 15
Atlantic puffin:H79
CANTUZ010000009.1:9790923
Log Adj R² = 0.317
n = 15
Atlantic puffin:H16
CANTUZ010000013.1:4114257
Log Adj R² = 0.241
n = 15
Atlantic puffin:H16
CANTUZ010000013.1:4114242
Log Adj R² = 0.237
n = 15
Atlantic puffin:H79
CANTUZ010000009.1:9790874
Log Adj R² = 0.219
n = 15
Atlantic puffin:H79
CANTUZ010000009.1:9790907
Log Adj R² = 0.214
n = 15
Atlantic puffin:H131
CANTUZ010000006.1:37349764
Log Adj R² = 0.191
n = 15
Atlantic puffin:H131
CANTUZ010000006.1:37349739
Log Adj R² = 0.184
n = 15
Atlantic puffin:H125b
CANTUZ010000002.1:93673315
Log Adj R² = 0.166
n = 15
Atlantic puffin:H131
CANTUZ010000006.1:37349682
Log Adj R² = 0.165
n = 15
Atlantic puffin:H79
CANTUZ010000009.1:9790872
Log Adj R² = 0.162
n = 15
Atlantic puffin:H16
CANTUZ010000013.1:4114260
Log Adj R² = 0.148
n = 15
Atlantic puffin:H42
CANTUZ010000003.1:44947877
Log Adj R² = 0.052
n = 15
Atlantic puffin:H32
CANTUZ010000002.1:150435670
Log Adj R² = 0.019
n = 15
Methylation
Observed Age (Years)

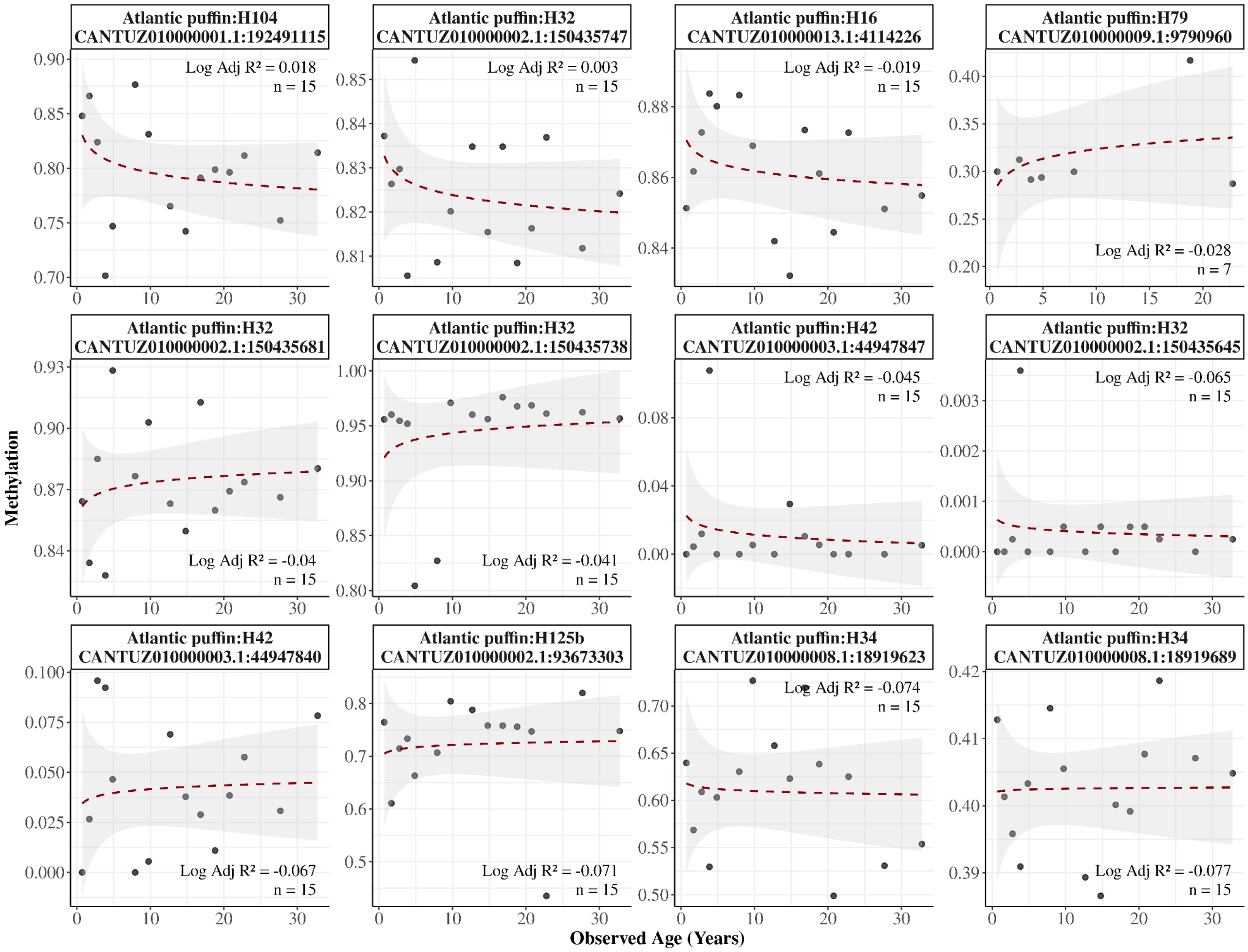


**Supplementary Figure 9. Age-dependent methylation in the Atlantic puffin blood samples**. Panels display methylation levels (y-axis) against observed age (x-axis) for each CpG site ($n_{Sample}$=15, $n_{CpG}$=32, $n_{Primer}$=10). The plot includes all detected CpG sites with more than 3 samples, excluding those in which more than half of the samples showed extreme methylation values (exactly 0 or 1). Annotations include primer number, genomic position, logarithmic adjusted $R^2$ (Log Adj $R^2$) between methylation and age, and number of samples. CpG sites are ordered by logarithmic adjusted $R^2$, and a logarithmic smoothing line is applied to each panel.

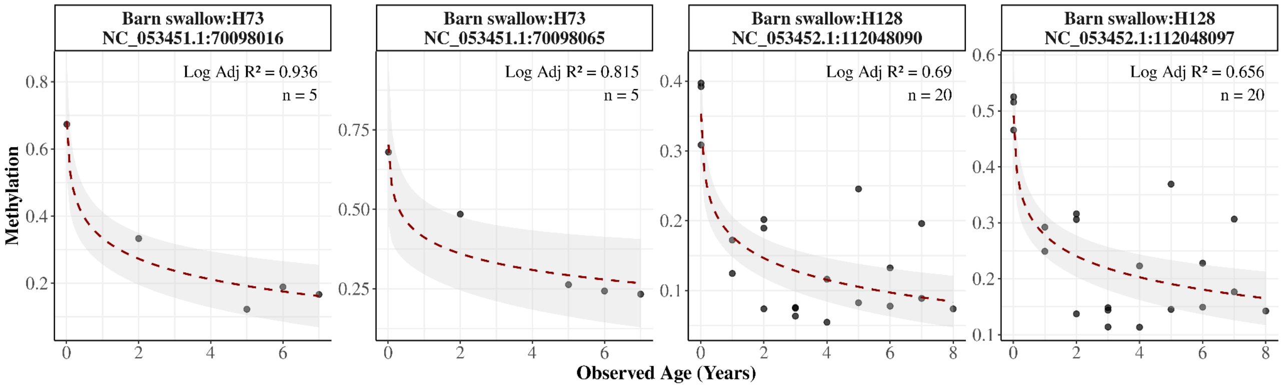


**Supplementary Figure 10. Age-dependent methylation in the Barn swallow blood samples**. Panels display methylation levels (y-axis) against observed age (x-axis) for each CpG site ($n_{Sample}$=20, $n_{CpG}$=4, $n_{Primer}$=2). The plot includes all detected CpG sites with more than 3 samples, excluding those in which more than half of the samples showed extreme methylation values (exactly 0 or 1). Annotations include primer number, genomic position, logarithmic adjusted $R^2$ (Log Adj $R^2$) between methylation and age, and number of samples. CpG sites are ordered by logarithmic adjusted $R^2$, and a logarithmic smoothing line is applied to each panel.

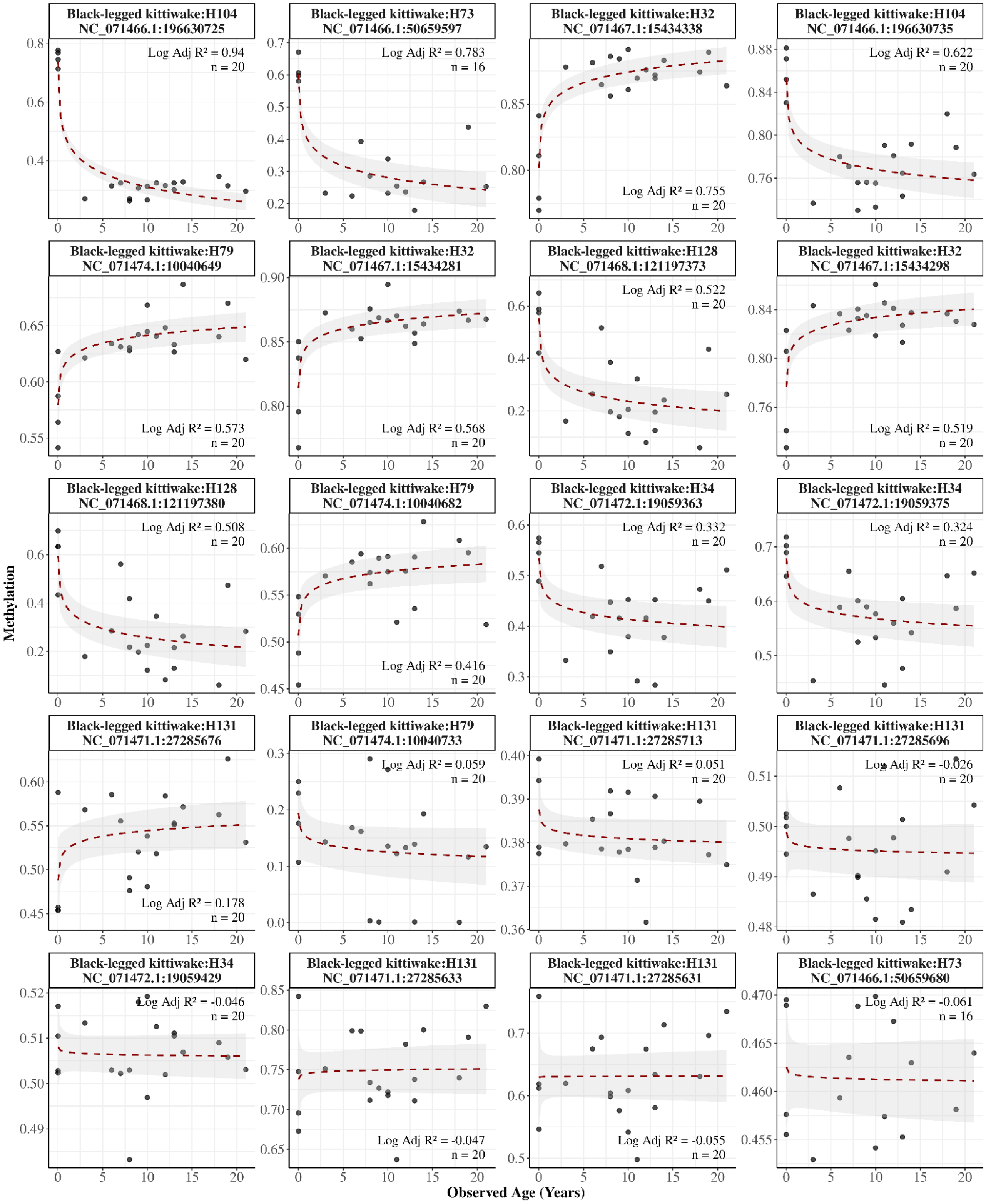


**Supplementary Figure 11. Age-dependent methylation in the Black-legged kittiwake blood samples**. Panels display methylation levels (y-axis) against observed age (x-axis) for each CpG site ($n_{Sample}$=20, $n_{CpG}$=20, $n_{Primer}$=7). The plot includes all detected CpG sites with more than 3 samples, excluding those in which more than half of the samples showed extreme methylation values (exactly 0 or 1). Annotations include primer number, genomic position, logarithmic adjusted $R^2$ (Log Adj $R^2$) between methylation and age, and number of samples. CpG sites are ordered by logarithmic adjusted $R^2$, and a logarithmic smoothing line is applied to each panel.

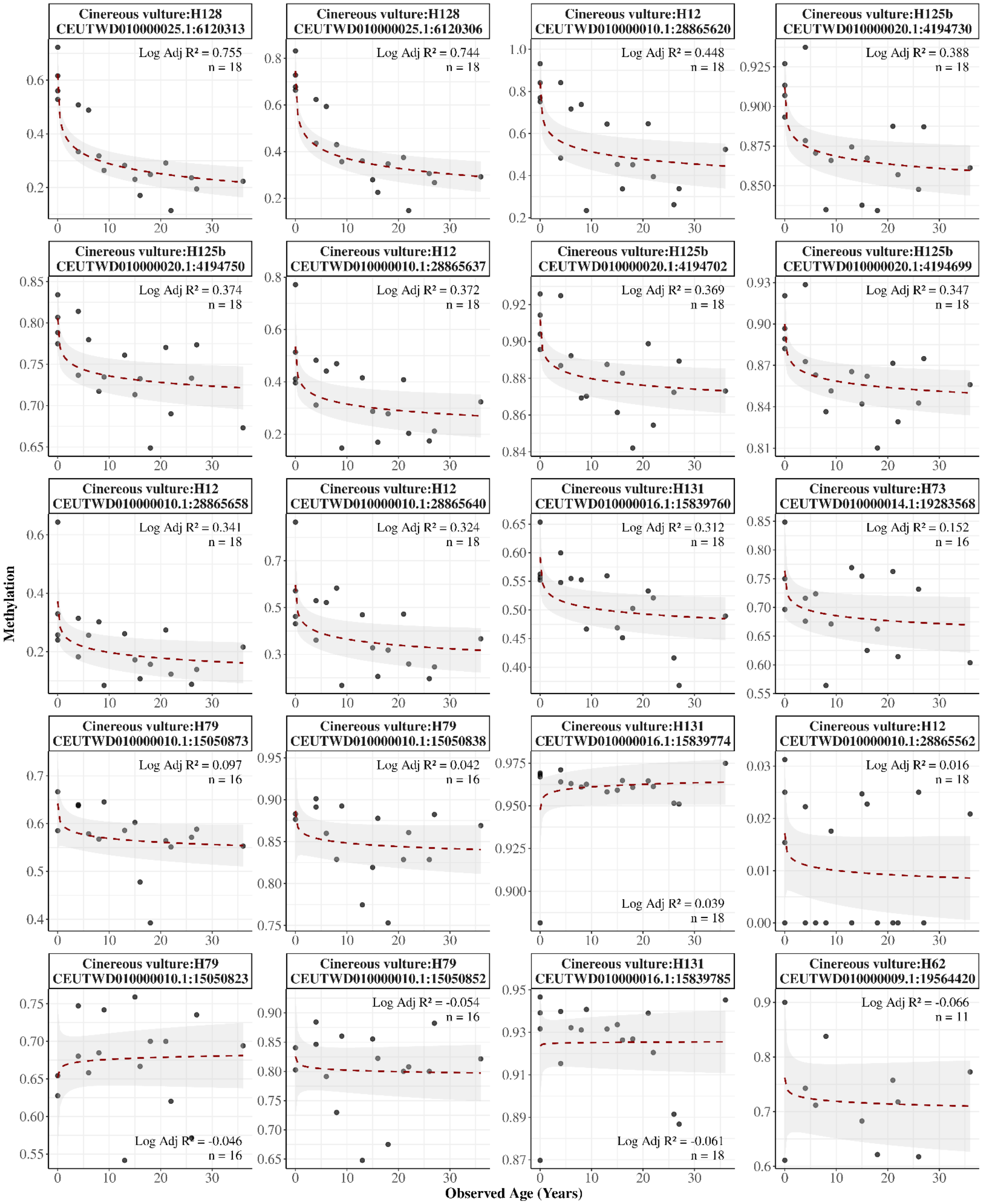

Cinereous vulture:H128 CEUTWD010000025.1:6120313
Log Adj R² = 0.755
n = 18
Cinereous vulture:H128 CEUTWD010000025.1:6120306
Log Adj R² = 0.744
n = 18
Cinereous vulture:H12 CEUTWD010000010.1:28865620
Log Adj R² = 0.448
n = 18
Cinereous vulture:H125b CEUTWD010000020.1:4194730
Log Adj R² = 0.388
n = 18
Cinereous vulture:H125b CEUTWD010000020.1:4194750
Log Adj R² = 0.374
n = 18
Cinereous vulture:H12 CEUTWD010000010.1:28865637
Log Adj R² = 0.372
n = 18
Cinereous vulture:H125b CEUTWD010000020.1:4194702
Log Adj R² = 0.369
n = 18
Cinereous vulture:H125b CEUTWD010000020.1:4194699
Log Adj R² = 0.347
n = 18
Cinereous vulture:H12 CEUTWD010000010.1:28865658
Log Adj R² = 0.341
n = 18
Cinereous vulture:H12 CEUTWD010000010.1:28865640
Log Adj R² = 0.324
n = 18
Cinereous vulture:H131 CEUTWD010000016.1:15839760
Log Adj R² = 0.312
n = 18
Cinereous vulture:H73 CEUTWD010000014.1:19283568
Log Adj R² = 0.152
n = 16
Cinereous vulture:H79 CEUTWD010000010.1:15050873
Log Adj R² = 0.097
n = 16
Cinereous vulture:H79 CEUTWD010000010.1:15050838
Log Adj R² = 0.042
n = 16
Cinereous vulture:H131 CEUTWD010000016.1:15839774
Log Adj R² = 0.039
n = 18
Cinereous vulture:H12 CEUTWD010000010.1:28865562
Log Adj R² = 0.016
n = 18
Cinereous vulture:H79 CEUTWD010000010.1:15050823
Log Adj R² = -0.046
n = 16
Cinereous vulture:H79 CEUTWD010000010.1:15050852
Log Adj R² = -0.054
n = 16
Cinereous vulture:H131 CEUTWD010000016.1:15839785
Log Adj R² = -0.061
n = 18
Cinereous vulture:H62 CEUTWD010000009.1:19564420
Log Adj R² = -0.066
n = 11
Methylation
Observed Age (Years)

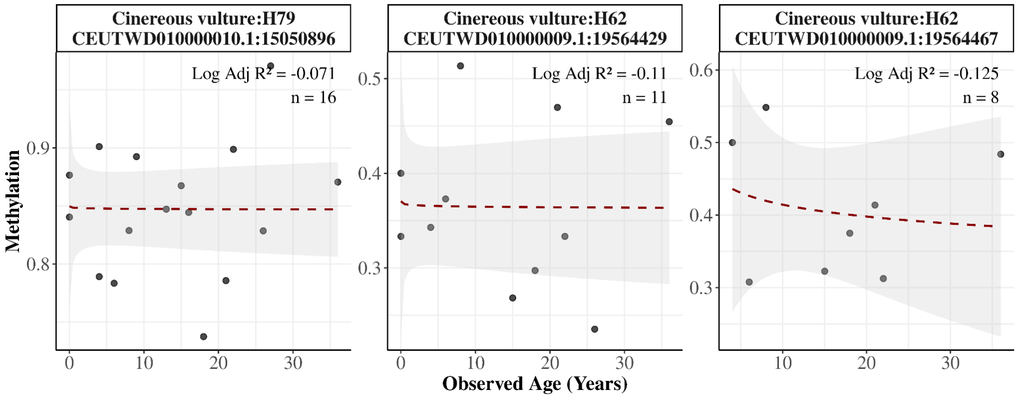


**Supplementary Figure 12. Age-dependent methylation in the Cinereous vulture blood samples**.Panels display methylation levels (y-axis) against observed age (x-axis) for each CpG site ($n_{Sample}$=18, $n_{CpG}$=23, $n_{Primer}$=7). The plot includes all detected CpG sites with more than 3 samples, excluding those in which more than half of the samples showed extreme methylation values (exactly 0 or 1). Annotations include primer number, genomic position, logarithmic adjusted $R^2$ (Log Adj $R^2$) between methylation and age, and number of samples. CpG sites are ordered by logarithmic adjusted $R^2$, and a logarithmic smoothing line is applied to each panel.

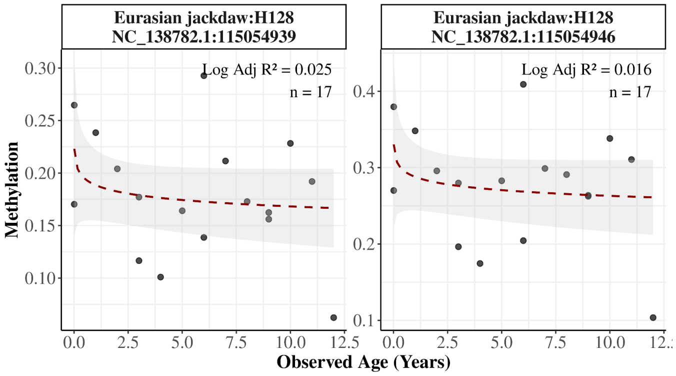


**Supplementary Figure 13. Age-dependent methylation in the Eurasian jackdaw blood samples**. Panels display methylation levels (y-axis) against observed age (x-axis) for each CpG site ($n_{Sample}$=17, $n_{CpG}$=2, $n_{Primer}$=1). The plot includes all detected CpG sites with more than 3 samples, excluding those in which more than half of the samples showed extreme methylation values (exactly 0 or 1). Annotations include primer number, genomic position, logarithmic adjusted $R^2$ (Log Adj $R^2$) between methylation and age, and number of samples. CpG sites are ordered by logarithmic adjusted $R^2$, and a logarithmic smoothing line is applied to each panel.

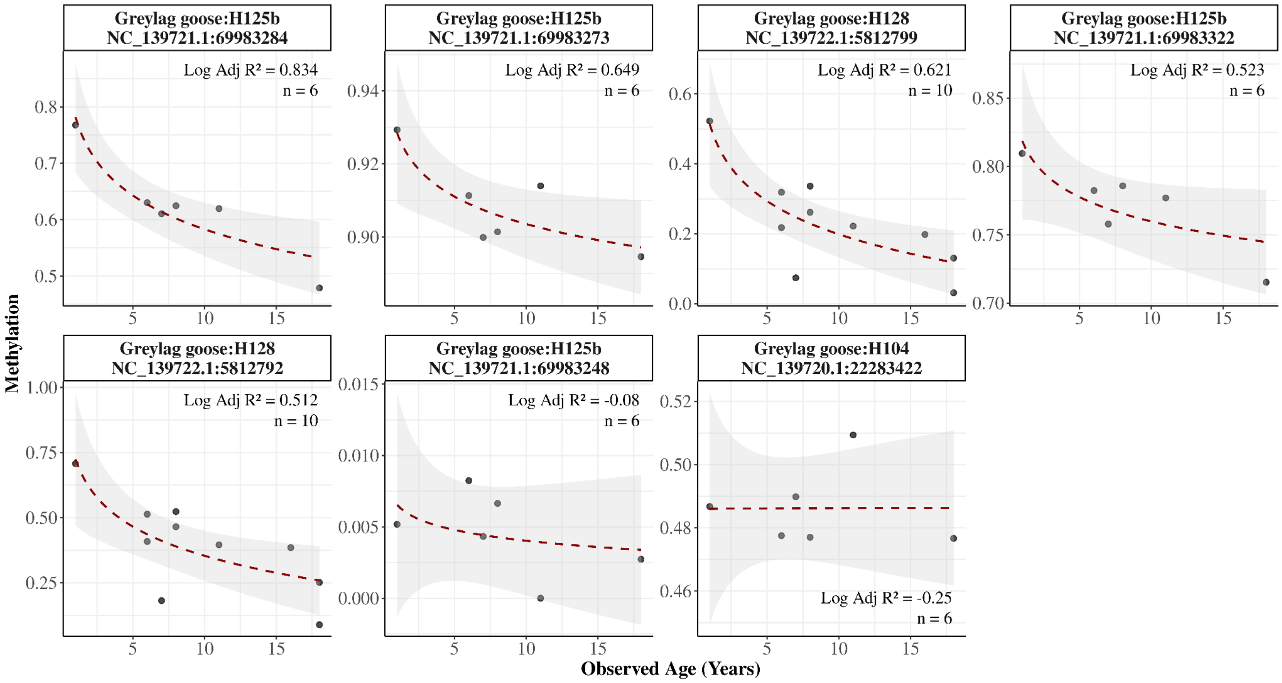


**Supplementary Figure 14. Age-dependent methylation in the Greylag goose blood samples**. Panels display methylation levels (y-axis) against observed age (x-axis) for each CpG site ($n_{Sample}$=10, $n_{CpG}$=7, $n_{Primer}$=3). The plot includes all detected CpG sites with more than 3 samples, excluding those in which more than half of the samples showed extreme methylation values (exactly 0 or 1). Annotations include primer number, genomic position, logarithmic adjusted $R^2$ (Log Adj $R^2$) between methylation and age, and number of samples. CpG sites are ordered by logarithmic adjusted $R^2$, and a logarithmic smoothing line is applied to each panel.

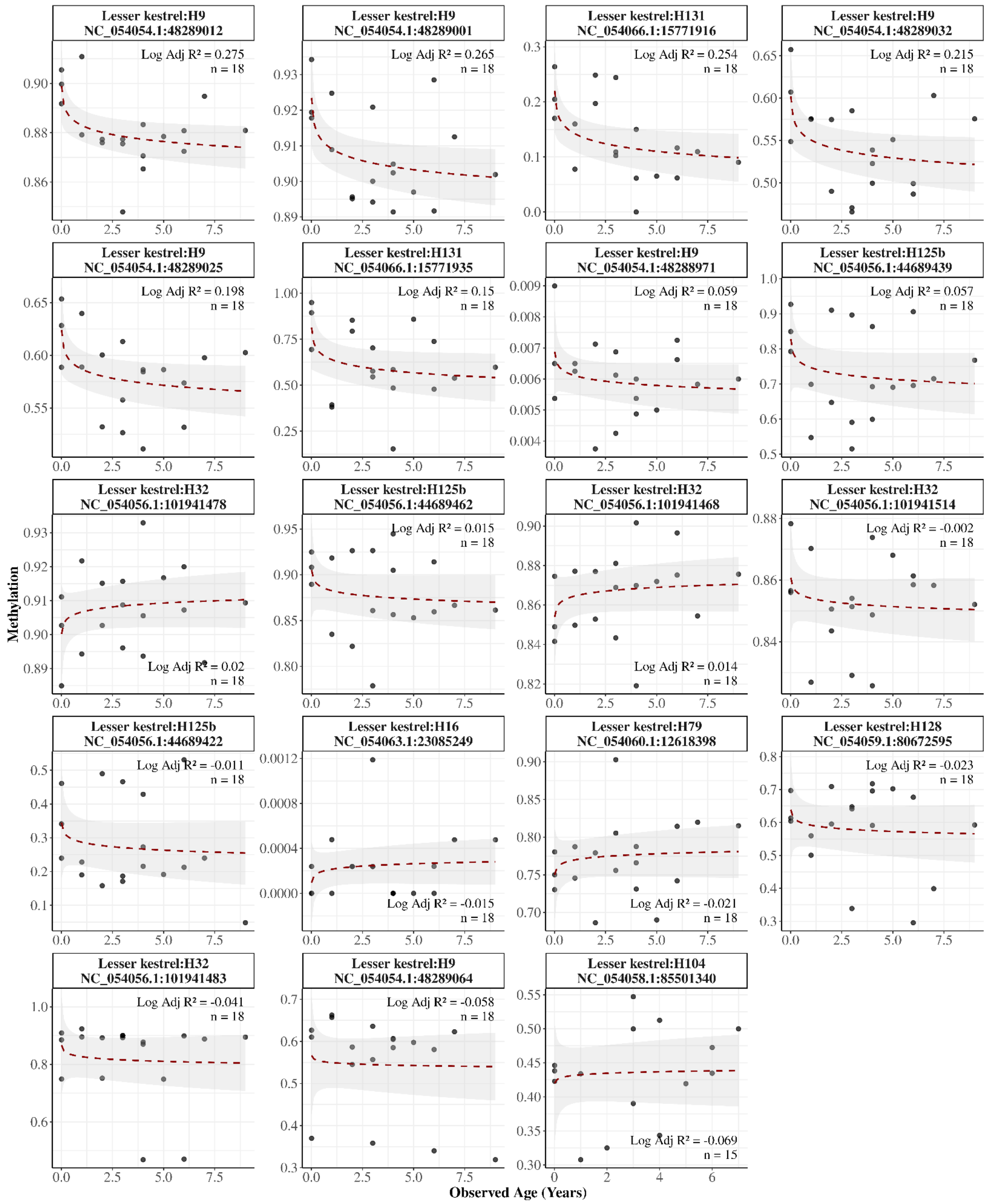


**Supplementary Figure 15. Age-dependent methylation in the Lesser kestrel blood samples**. Panels display methylation levels (y-axis) against observed age (x-axis) for each CpG site ($n_{Sample}$=18, $n_{CpG}$=19, $n_{Primer}$=8). The plot includes all detected CpG sites with more than 3 samples, excluding those in which more than half of the samples showed extreme methylation values (exactly 0 or 1).

Annotations include primer number, genomic position, logarithmic adjusted $R^2$ (Log Adj $R^2$) between methylation and age, and number of samples. CpG sites are ordered by logarithmic adjusted $R^2$, and a logarithmic smoothing line is applied to each panel.

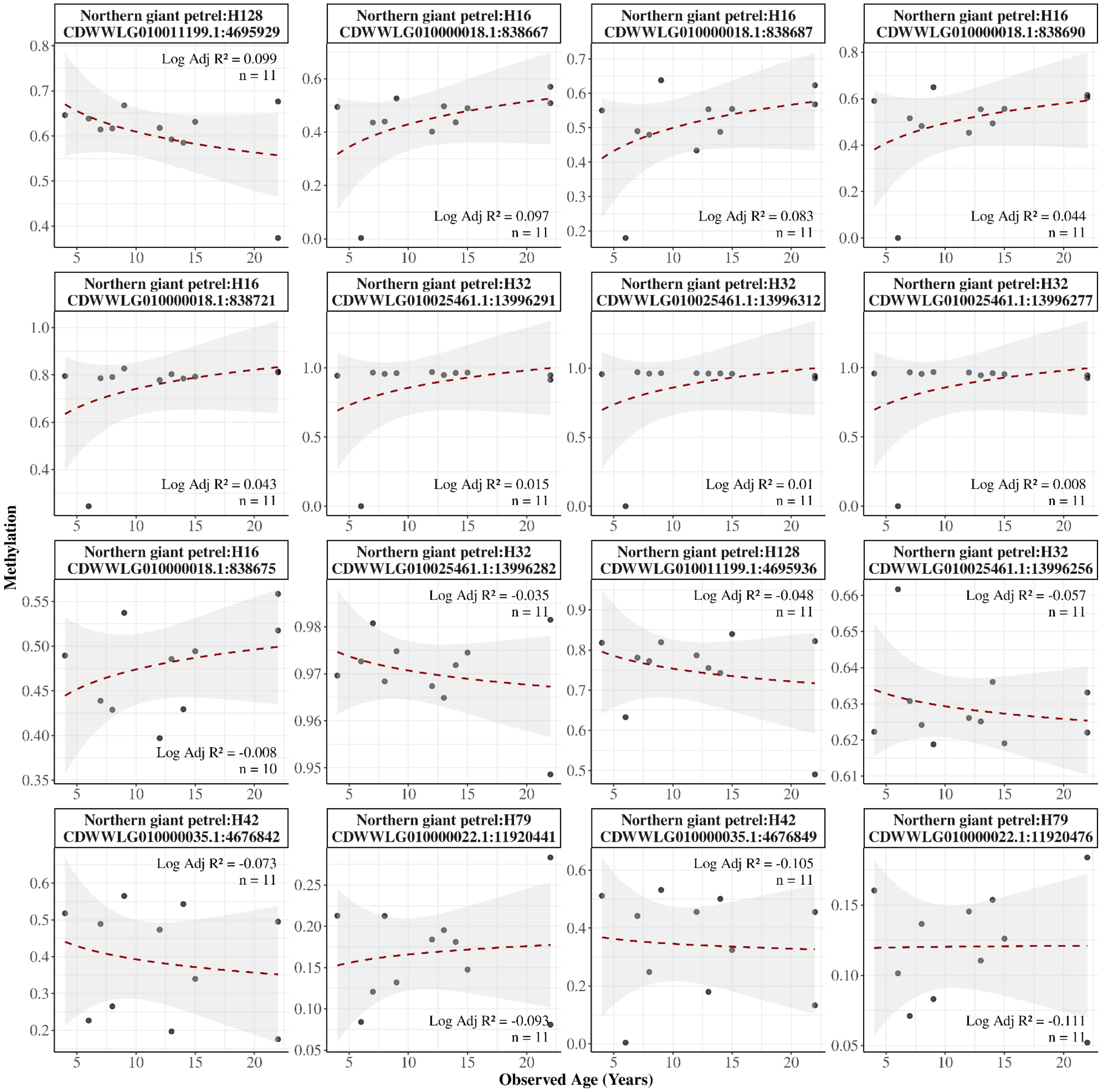


**Supplementary Figure 16. Age-dependent methylation in the Northern giant petrel muscle samples**. Panels display methylation levels (y-axis) against observed age (x-axis) for each CpG site ($n_{Sample}$=11, $n_{CpG}$=16, $n_{Primer}$=5). The plot includes all detected CpG sites with more than 3 samples, excluding those in which more than half of the samples showed extreme methylation values (exactly 0 or 1). Annotations include primer number, genomic position, logarithmic adjusted $R^2$ (Log Adj $R^2$) between methylation and age, and number of samples. CpG sites are ordered by logarithmic adjusted $R^2$, and a logarithmic smoothing line is applied to each panel.

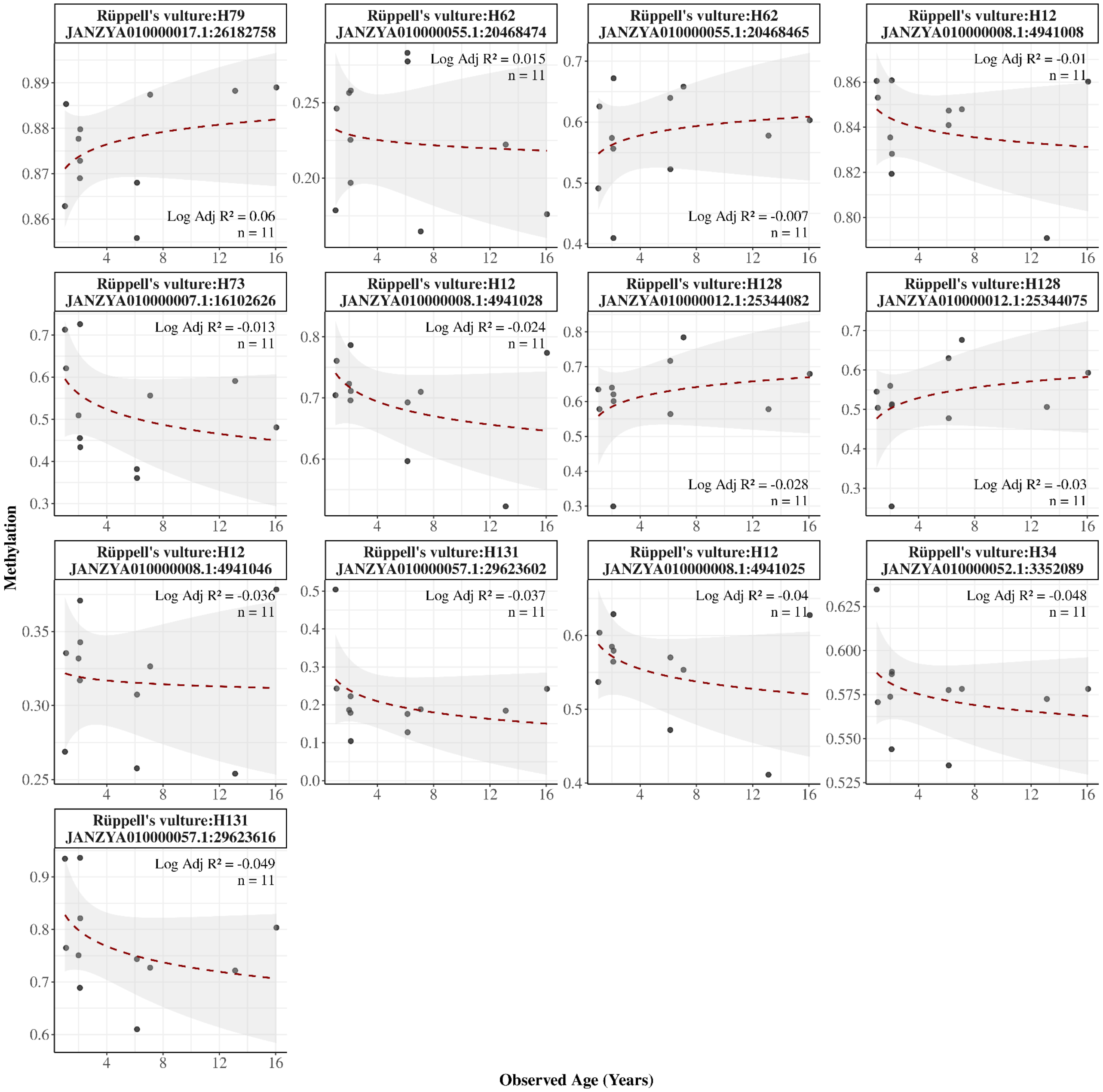


**Supplementary Figure 17. Age-dependent methylation in the Rüppell's vulture blood samples**. Panels display methylation levels (y-axis) against observed age (x-axis) for each CpG site ($n_{Sample}$=11, $n_{CpG}$=13, $n_{Primer}$=7). The plot includes all detected CpG sites with more than 3 samples, excluding those in which more than half of the samples showed extreme methylation values (exactly 0 or 1). Annotations include primer number, genomic position, logarithmic adjusted $R^2$ (Log Adj $R^2$) between methylation and age, and number of samples. CpG sites are ordered by logarithmic adjusted $R^2$, and a logarithmic smoothing line is applied to each panel.

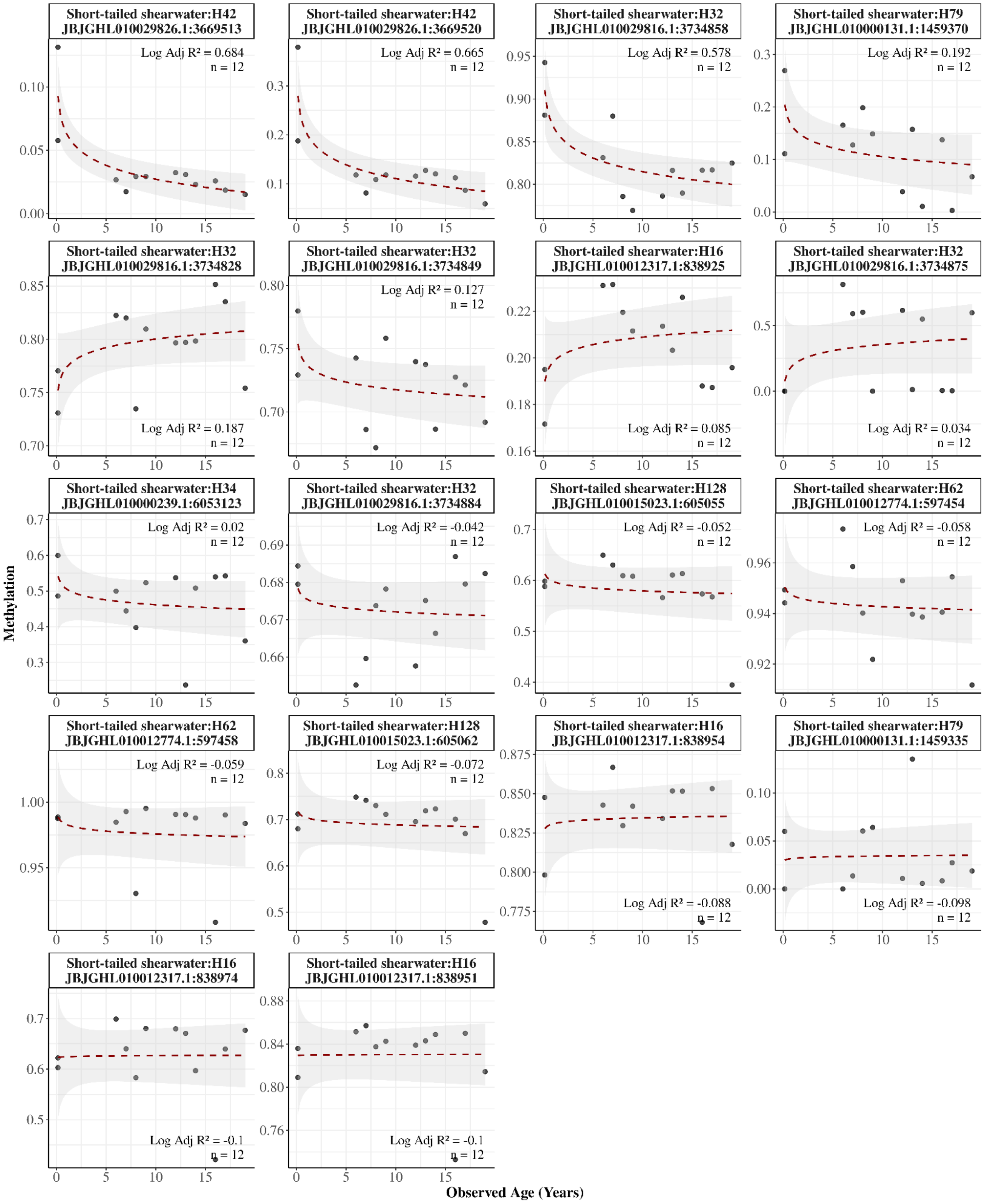


**Supplementary Figure 18. Age-dependent methylation in the Short-tailed shearwater blood samples**. Panels display methylation levels (y-axis) against observed age (x-axis) for each CpG site ($n_{Sample}$=12, $n_{CpG}$=18, $n_{Primer}$=7). The plot includes all detected CpG sites with more than 3 samples, excluding those in which more than half of the samples showed extreme methylation values (exactly 0 or 1). Annotations include primer number, genomic position, logarithmic adjusted $R^2$ (Log Adj $R^2$) between methylation and age, and number of samples. CpG sites are ordered by logarithmic adjusted $R^2$, and a logarithmic smoothing line is applied to each panel.

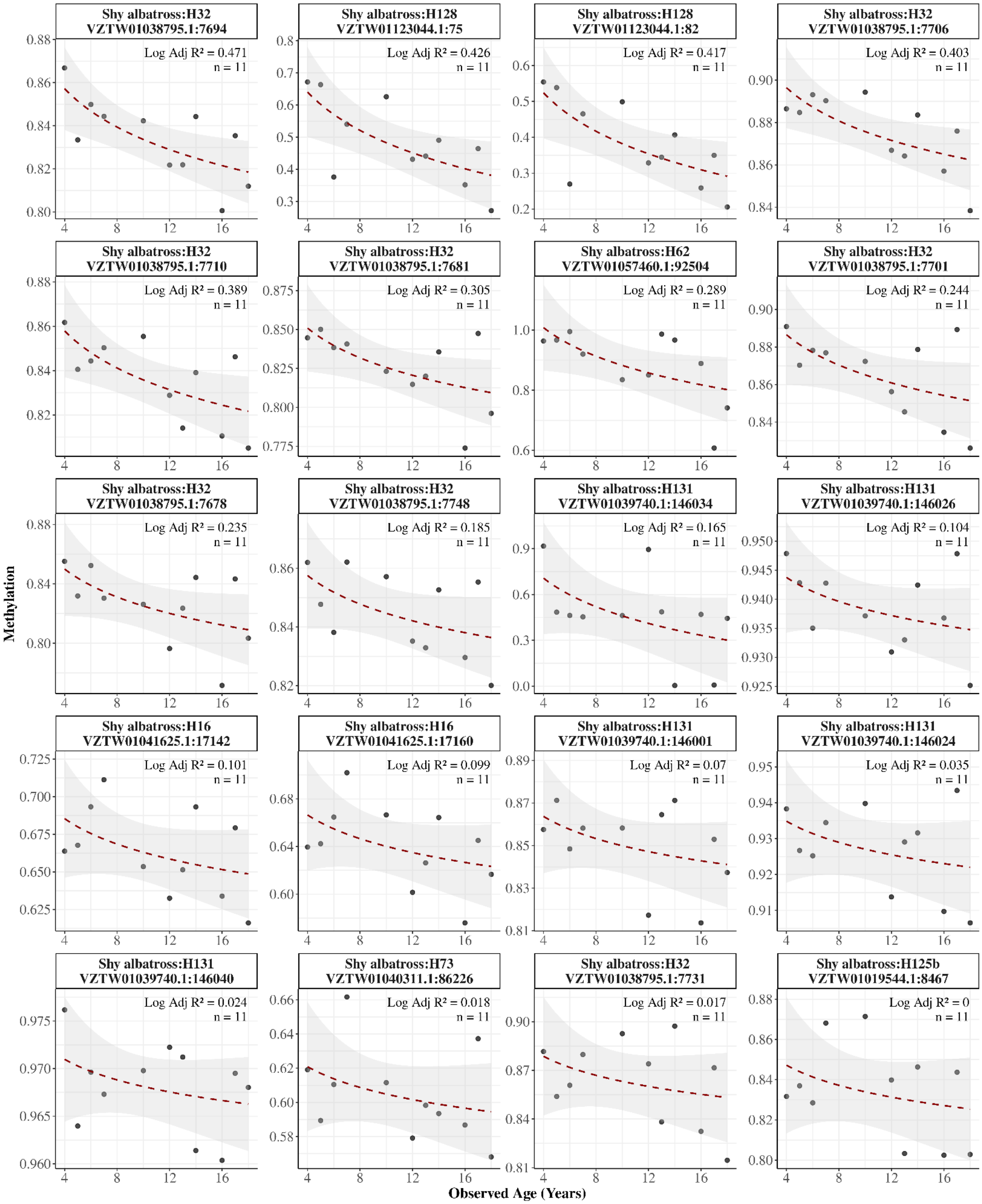

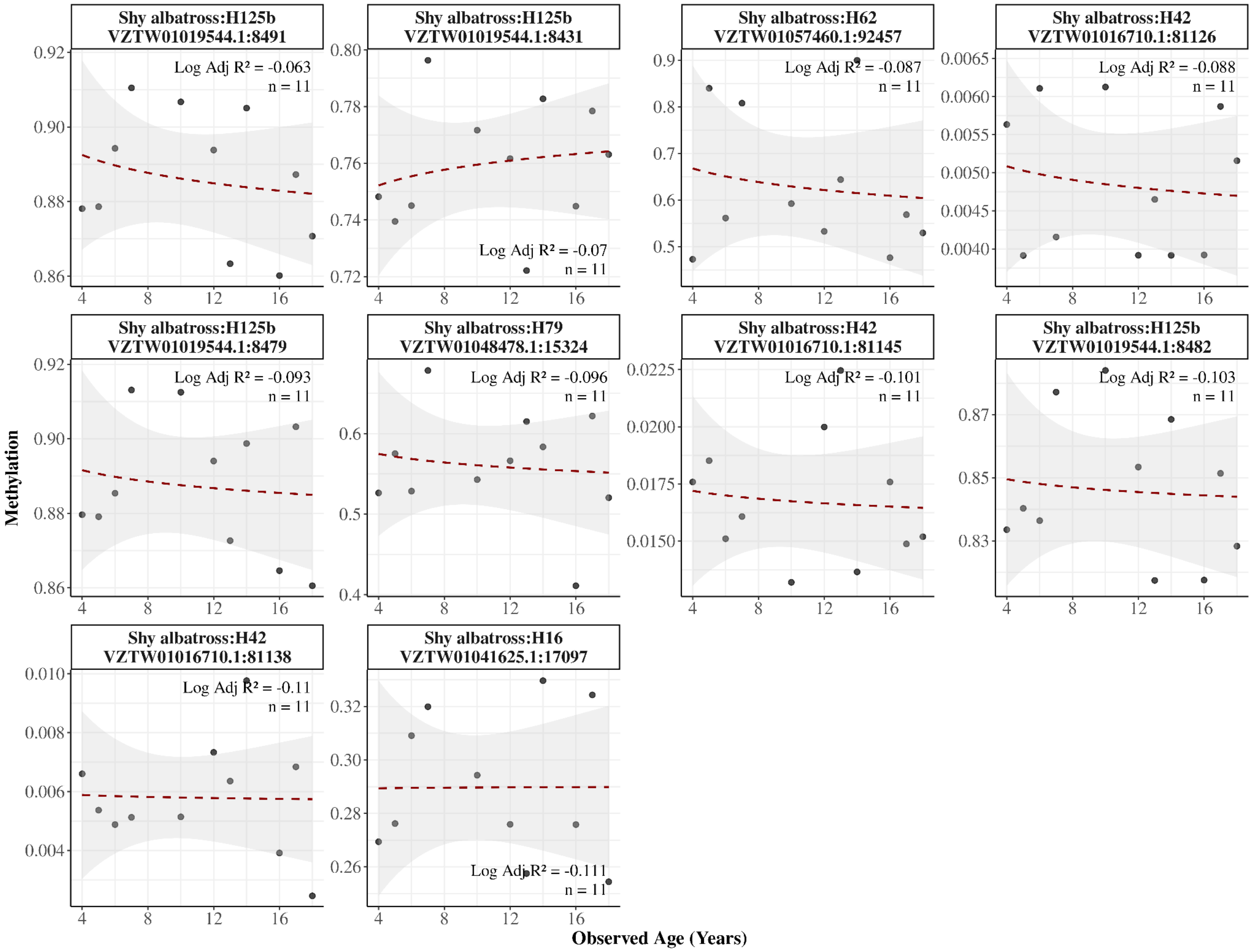


**Supplementary Figure 19. Age-dependent methylation in the Shy albatross blood samples**. Panels display methylation levels (y-axis) against observed age (x-axis) for each CpG site ($n_{Sample}$=11, $n_{CpG}$=30, $n_{Primer}$=9). The plot includes all detected CpG sites with more than 3 samples, excluding those in which more than half of the samples showed extreme methylation values (exactly 0 or 1). Annotations include primer number, genomic position, logarithmic adjusted $R^2$ (Log Adj $R^2$) between methylation and age, and number of samples. CpG sites are ordered by logarithmic adjusted $R^2$, and a logarithmic smoothing line is applied to each panel.

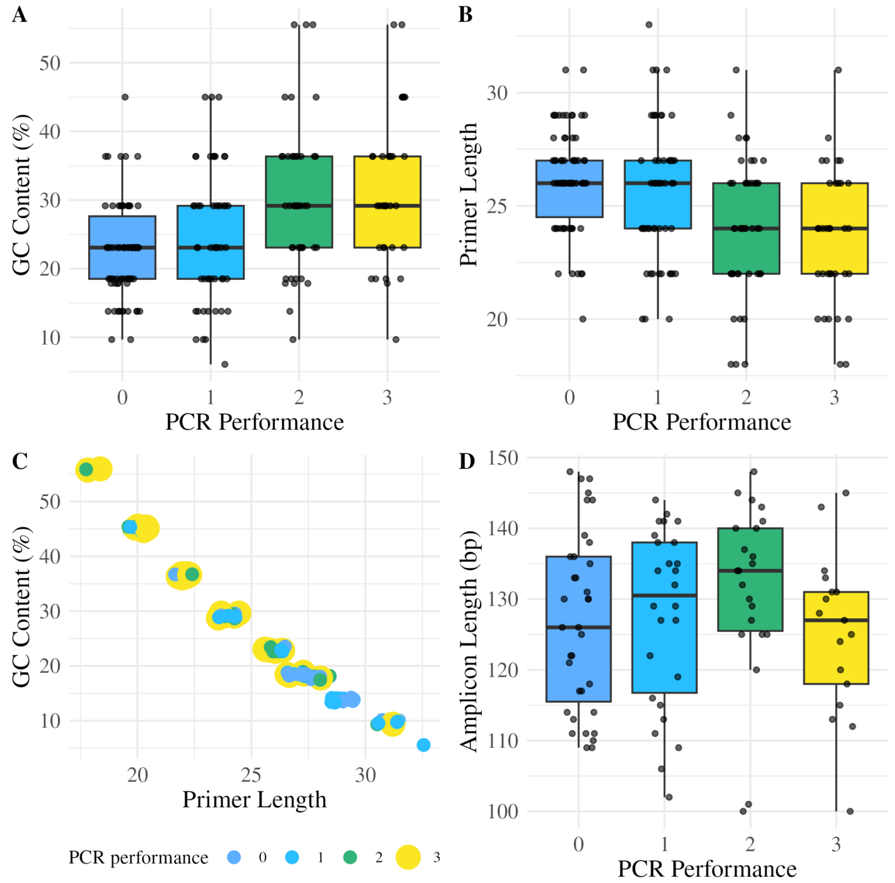


**Supplementary Figure 20. PCR performance of individually tested primer pairs (n=100).** Primer pairs were screened individually using King penguin BS-DNA pooled from 10 individuals, and the performance was determined by gel electrophoresis. PCR performance is shown in relation to GC content (A), primer length in bases (B). Panel D shows the relationship between GC content and primer length, while panel D illustrates the relationship between PCR performance and amplicon length in base pairs (bp).

### Supplementary Data

**Supplementary Data 1. Suggested primer versions for different species.** Alternative versions of BEAC primers (rows) are given for each species (columns) tested in the manuscript. Submitted as a separate file. Screenshot below for illustration.

**Supplementary Data 2. Fasta-file of flanking sequences of the BEAC panel.** Sequences used in primer design of the 24 BEAC primer pairs, 200 bases up- and downstream of the target loci. Screenshot below for illustration.

**Supplementary Data 3. Scripts used in BEAC development.** R scripts used in the BEAC model training
